# A computational screen for alternative genetic codes in over 250,000 genomes

**DOI:** 10.1101/2021.06.18.448887

**Authors:** Yekaterina Shulgina, Sean R. Eddy

## Abstract

The genetic code has been proposed to be a “frozen accident”, but the discovery of alternative genetic codes over the past four decades has shown that it can evolve to some degree. Since most examples were found anecdotally, it is difficult to draw general conclusions about the evolutionary trajectories of codon reassignment and why some codons are affected more frequently. To fill in the diversity of genetic codes, we developed Codetta, a computational method to predict the amino acid decoding of each codon from nucleotide sequence data. We surveyed the genetic code usage of over 250,000 bacterial and archaeal genome sequences in GenBank and discovered five new reassignments of arginine codons (AGG, CGA, and CGG), representing the first sense codon changes in bacteria. In a clade of uncultivated Bacilli, the reassignment of AGG to become the dominant methionine codon likely evolved by a change in the amino acid charging of an arginine tRNA. The reassignments of CGA and/or CGG were found in genomes with low GC content, an evolutionary force which likely helped drive these codons to low frequency and enable their reassignment.

## Introduction

The genetic code defines how mRNA sequences are decoded into proteins. The ancient origin of the standard genetic code is reflected in its near-universal usage, once proposed to be a “frozen accident” that is too integral to the translation of all proteins to change (***Crick, 1968***). However, the discovery of alternative genetic codes in over 30 different lineages of bacteria, eukaryotes, and mitochondria over the past four decades has made it clear that the genetic code is capable of evolving to some degree (***Knight et al., 2001a***; ***Kollmar and Mühlhausen, 2017***).

The first alternative genetic codes were discovered by comparing newly sequenced genomes to amino acid sequences obtained by direct protein sequencing. Nonstandard codon translations were found this way in human mitochondria (***Barrell et al., 1979***), *Candida* yeasts (***Kawaguchi et al., 1989***), green algae (***Schneider et al., 1989***), and *Euplotes* ciliates (***Meyer et al., 1991***). Some reassignments of stop codons to amino acids were detected from DNA sequence alone, based on the appearance of in-frame stop codons in critical genes (***Yamao et al., 1985***; ***Caron and Meyer, 1985***; ***Cupples and Pearlman, 1986***; ***Keeling and Doolittle, 1996***; ***McCutcheon et al., 2009***; ***Campbell et al., 2013***; ***Záhonová et al., 2016***). As DNA sequence data has accumulated faster than direct protein sequences, computational methods have been developed to predict the genetic code from DNA sequence. The core principle of most methods is to align genomic coding regions to homologous sequences in other organisms (creating multiple sequence alignments) and then totally the most frequent amino acid aligned to each of the 64 codons. This approach led to the discovery of new genetic codes in screens of ciliates (***Swart et al., 2016***; ***Heaphy et al., 2016***), yeasts (***Riley et al., 2016***; ***Krassowski et al., 2018***), green algal mitochondria (***Noutahi et al., 2019***; ***Žihala and Eliáš, 2019***) and stop codon reassignments in metagenomic data (***Ivanova et al., 2014***) and the development of software for specific phylogenetic groups (***Abascal et al., 2006b***; ***Mühlhausen and Kollmar, 2014***; ***Noutahi et al., 2017***). Some approaches, such as FACIL (***Dutilh et al., 2011***), have expanded phylogenetic breadth by using profile Hidden Markov model (HMM) representations of conserved proteins from phylogenetically diverse databases such as Pfam (***El-Gebali et al., 2019***). However, a systematic survey of genetic code usage across the tree of life has not yet been possible. Existing methods are generally either 1) phylogenetically restricted to clades where multiple sequence alignments can be built for a predetermined set of proteins or 2) lacking sufficiently robust and objective statistical footing to enable a large-scale screen with high accuracy.

A potentially incomplete set of alternative genetic codes limits our ability to understand the evolutionary processes behind codon reassignment. One open question is why some codon reassignments reappear independently. Reassignment of the stop codons UAA and UAG to glutamine is the most common change in eukaryotic nuclear genomes, appearing at least five independent times (***Schneider et al., 1989***; ***Keeling and Doolittle, 1996***; ***Keeling and Leander, 2003***; ***Karpov et al., 2013***; ***Swart et al., 2016***). In bacteria, all of the known changes reassign the stop codon UGA to either glycine in the Absconditabacteria and Gracilibacteria (***Campbell et al., 2013***; ***Rinke et al., 2013***) or tryptophan in the Mycoplasmatales, Entomoplasmatales (***Bové, 1993***), and several insect endosymbiotic bacteria (***McCutcheon et al., 2009***; ***McCutcheon and Moran, 2010***; ***Bennett and Moran, 2013***; ***Salem et al., 2017***). These recurring changes may reflect which codon reassignments are easier to evolve due to pre-existing constraints on tRNA anticodon-codon pairing, aminoacyl-tRNA synthetase recognition of cognate tRNAs, release factor binding, and other key steps in translation. However, without a complete picture of genetic code diversity, it is hard to disentangle patterns of codon reassignment from observation bias. For instance, in-frame stop codons caused by a stop codon reassignment may be more easily detectable than a subtle change in amino acid conservation indicative of a sense codon reassignment.

Another open question is how a new codon meaning can evolve without disrupting the translation of most proteins. Reassigning a codon leads to the incorporation of the incorrect amino acid at all preexisting codon positions (***Crick, 1968***). Three evolutionary models differ in the pressure driving substitutions to remove the codon from positions that cannot tolerate the new translation. In the ‘codon capture’ model, the codon is first driven to near-extinction by pressures unrelated to reassignment, such as biased genomic GC content or genome reduction, which then minimizes the impact of reassignment on protein translation (***Osawa and Jukes, 1989***). This model was first proposed for the reassignment of the stop codon UGA to tryptophan in *Mycoplasma capricolum*, whose low genomic GC content (25% GC) in combination with small genome size (1 Mb) was thought to have driven the stop codon UGA to extremely low usage in favor of UAA and allowed ‘capture’ of UGA by a tryptophan tRNA (***Bové, 1993***; ***Osawa and Jukes, 1989***). For larger nuclear genomes, other models have been proposed where codon usage changes occur concurrently with, and are driven by, changes in decoding capability. In the ‘ambiguous intermediate’ model, a codon is decoded stochastically as two different meanings in an intermediate step of codon reassignment, and this translational pressure induces codon substitutions at positions where ambiguity is deleterious (***Schultz and Yarus, 1994***; ***Massey et al., 2003***). Extant examples of ambiguous translation support the plausibility of this model, such as yeasts that translate the codon CUG as both leucine and serine by stochastic tRNA charging (***Gomes et al., 2007***) or by competing tRNA species (***Mühlhausen et al., 2018***). Alternatively, the ‘tRNA loss driven reassignment’ model proposes an intermediate stage where a codon cannot be translated efficiently, perhaps due to tRNA gene loss or mutation, creating pressure for synonymous substitutions specifically away from that codon, allowing it to be captured later by a different tRNA (***Mühlhausen et al., 2016***; ***Sengupta and Higgs, 2005***). These three models are not mutually exclusive and substitutions at the reassigned codon can occur due to a combination of these pressures.

Here, we describe Codetta, a computational method for predicting the genetic code which can scale to analyze thousands of genomes. We perform the first survey of genetic code usage in all bacterial and archaeal genomes, reidentifying all known codes in the dataset and discovering the first examples of sense codon changes in bacteria. All five reassignments affect arginine codons (AGG, CGA, and CGG) and provide clues to help us understand how alternative genetic codes evolve.

## Results

### Codetta: a computational method to infer the genetic code

We developed Codetta, a computational method that takes DNA or RNA sequences from a single organism and predicts an amino acid translation for each of the 64 codons. The general idea is to align the input nucleotide sequence to probabilistic profiles of conserved protein domains in order to obtain, for each of the 64 codons, a set of profile positions aligned to that codon. Each profile position has twenty probabilities describing the expected amino acid. For each of the 64 codons, we aggregate over the set of aligned profile positions to infer the single most likely amino acid decoding of the codon. Most previous approaches for genetic code prediction use the same basic idea (***Abascal et al., 2006b***; ***Dutilh et al., 2011***; ***Mühlhausen and Kollmar, 2014***; ***Swart et al., 2016***; ***Heaphy et al., 2016***; ***Riley et al., 2016***; ***Krassowski et al., 2018***; ***Noutahi et al., 2019***), typically aligning the input sequence to multiple sequence alignments and using a simple rule to select the best amino acid for each codon.

With Codetta, we extend this idea to systematic high-throughput analysis by using a probabilistic modeling approach to infer codon decodings, and by taking advantage of the large collection of probabilistic profiles of conserved protein domains (profile HMMs) in the Pfam database (***El-Gebali et al., 2019***). Profile HMMs are built from multiple sequence alignments, and the emission probabilities at each consensus column are estimates of the expected amino acid frequencies. The Pfam database contains over 17,000 profile HMMs of conserved protein domains from all three domains of life, which are expected to align to about 50% of coding regions in a genome (***El-Gebali et al., 2019***). We align Pfam profile HMMs to a six-frame standard genetic code translation of the input DNA/RNA sequence using the HMMER `hmmscan` program (***Figure 1A***). Since we rely on a preliminary standard code translation, conserved protein domains could fail to align in organisms using radically different genetic codes. In the set of statistically significant `hmmscan` alignments (E-value < 10^−10^), we make the simplifying approximation of considering each aligned consensus column independently, so the alignments are viewed as a set of pairwise associations between a codon *z* (64 possibilities) and a consensus column of a Pfam domain profile (denoted *C*, an index identifying a Pfam consensus column).

**Figure 1.**
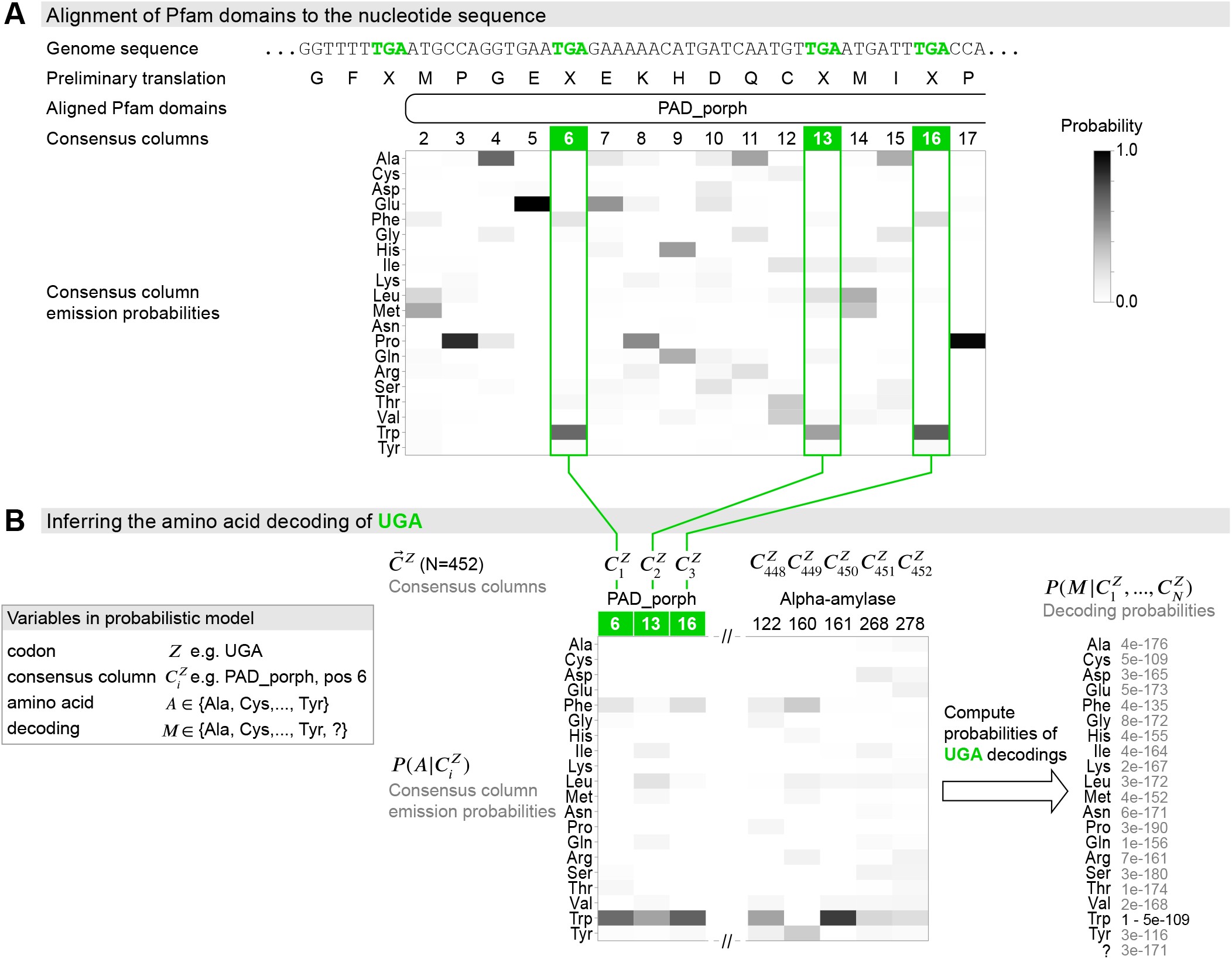
Schematic of the genetic code inference method implemented in Codetta. (A) A fragment of the *Mycoplasma capricolum* genome is used to demonstrate alignment of a Pfam domain (PAD_porph) to a preliminary standard code translation of the input DNA sequence (one of six frames shown). All canonical stop codons, including UGA (TGA in genome sequence, reassigned to tryptophan in *M. capricolum*), are translated as ‘X’ in the preliminary standard code translation which `hmmscan` (program used to align Pfam domains) treats as an unknown amino acid. Each consensus column in the PAD_porph domain has a characteristic emission probability for each of the twenty canonical amino acids, represented by a heatmap. (B) Pfam consensus columns aligning to UGA codons across the entire genome comprise the 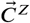 set for UGA (*N*=452 Pfam consensus columns). The Pfam emission probabilities 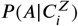 for all 452 aligned consensus columns are used to compute the decoding probabilities 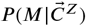. The most likely amino acid translation of UGA is inferred to be tryptophan, with decoding probability greater than the cutoff of 0.9999.

From these data, we infer each of the 64 codons one at a time (***Figure 1B***). For a codon *z* (e.g. UGA), the observed data 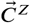 are a set of *N* consensus columns 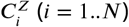 that associate to *z* in the provisional alignments. We model the main data-generative process abstractly, imagining that each column 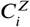 was drawn from the pool of all possible consensus columns by codon *z* which is translated as an unknown amino acid *A*. Each column has an affinity for codon *z* proportional to the column’s emission probability for the amino acid *A*, *P* (*A | C*). A consensus column strongly conserved for a particular amino acid *A* will tend to only associate with codons that translate to *A*; moreover, consensus columns weakly conserved for *A* may also associate with probability proportional to their conservation for *A*. Thus this abstract matching process generates an observed 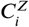 column association with the codon *z* (translated as amino acid *A*) with probability

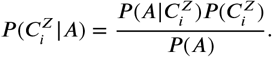

Here 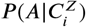 is the emission probability for amino acid *A* at the Pfam consensus column 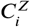. *P*(*A*) is the average emission probability for amino acid *A* over the pool of all possible consensus columns *C*, which we take to be all columns aligned to the target genome in order to better reflect genome-specific biases in amino acid usage.

Given the data 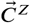 and this abstract generative model, we infer the most likely decoding *M* for codon *z* out of 21 possibilities *M* ∈ {Ala, Cys, …, Tyr, ?} (***Figure 1B***). The *M* = ? model of non-specific translation draws columns randomly and serves to catch codons that do not encode a specific amino acid, such as stop codons and ambiguously translated codons. For a given decoding *M*, the probability of the observed columns 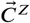 is then:

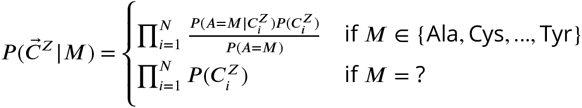

Setting the prior probability of each decoding, *P*(*M*), to be uniform, we compute the probability of the decoding *M* as:

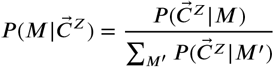

We assign an amino acid translation to a codon if it attains a decoding probability above some threshold (typically 0.9999). We assign a ‘?’ if no amino acid decoding satisfies the probability threshold (including the case where ‘?’ itself has high probability). A ‘?’ assignment tends to happen if the codon is rare, with few aligned Pfam consensus columns on which to base the inference, or if the codon is ambiguously translated such that no single amino acid model reaches high probability. Because we do not model stop codons explicitly, we expect ‘?’ to be the inferred meaning since stop codons ideally would have few or no aligned Pfam consensus columns.

To assess how many columns in 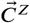 are needed for reliable codon assignment, we constructed synthetic 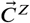 datasets ranging from 2 to 500 consensus columns by subsampling the consensus columns aligned to each of the 61 sense codons in the *Escherichia coli* genome. We calculated the per-codon error rate (fraction of samples predicting the incorrect amino acid) and the per-codon power (fraction of samples predicting the correct amino acid) using a probability threshold of 0.9999. Lack of an amino acid inference (‘?’) contributed to neither. Per-codon error rates were < 0.00005 for all sizes of 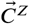 and we found that about 20 aligned consensus columns in 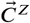 suffice for >95% power. Accuracy may differ in real genomes for various biological reasons, but these results gave us confidence to proceed.

### Genetic code prediction of 462 yeast species confirms known distributions of CUG reassignment

We further validated Codetta on the budding yeasts (Saccharomycetes, 462 sequenced species) which vary in their translation of CUG as either serine, leucine, or alanine depending on the species (***Mühlhausen et al., 2016***; ***Krassowski et al., 2018***; ***Mühlhausen et al., 2018***). In some CUG-Ser clade species, such as *Candida albicans*, CUG codons are stochastically decoded as a mix of serine (97%) and leucine (3%) because the CUG-decoding tRNA_CAG_ is aminoacylated by both the seryl- and leucyl-tRNA synthetases (***Suzuki et al., 1997***; ***Gomes et al., 2007***). Codetta is not designed to predict ambiguous decoding and is expected to assign either the dominant amino acid or a ‘?’ in cases like *C. albicans*.

For 453 species, the predicted CUG translation was consistent with the known phylogenetic distribution of CUG reassignments (***Figure 2A***). This includes *C. albicans*, which was predicted to use the predominant serine translation (***Gomes et al., 2007***). For the remaining nine species, Codetta did not put a high probability on any amino acid decoding of CUG (inferred meaning of ‘?’). Two of these species– *Babjeviella inositovora* and *Cephaloascus fragrans*– are basal members of the CUG-Ser clade. Both of these genomes contain a CUG-decoding tRNA_CAG_ gene with features of serine identity (see Methods) and *B. inositovora* has previously been shown to translate CUG codons primarily as serine by whole proteome mass spectrometry (***Krassowski et al., 2018***; ***Mühlhausen et al., 2018***), suggesting that CUG is decoded as serine in these species. Codetta did not infer an amino acid for CUG because the aligned Pfam consensus columns were not consistently conserved for a single amino acid (***Figure 2–Figure Supplement 1***).

**Figure 2.**
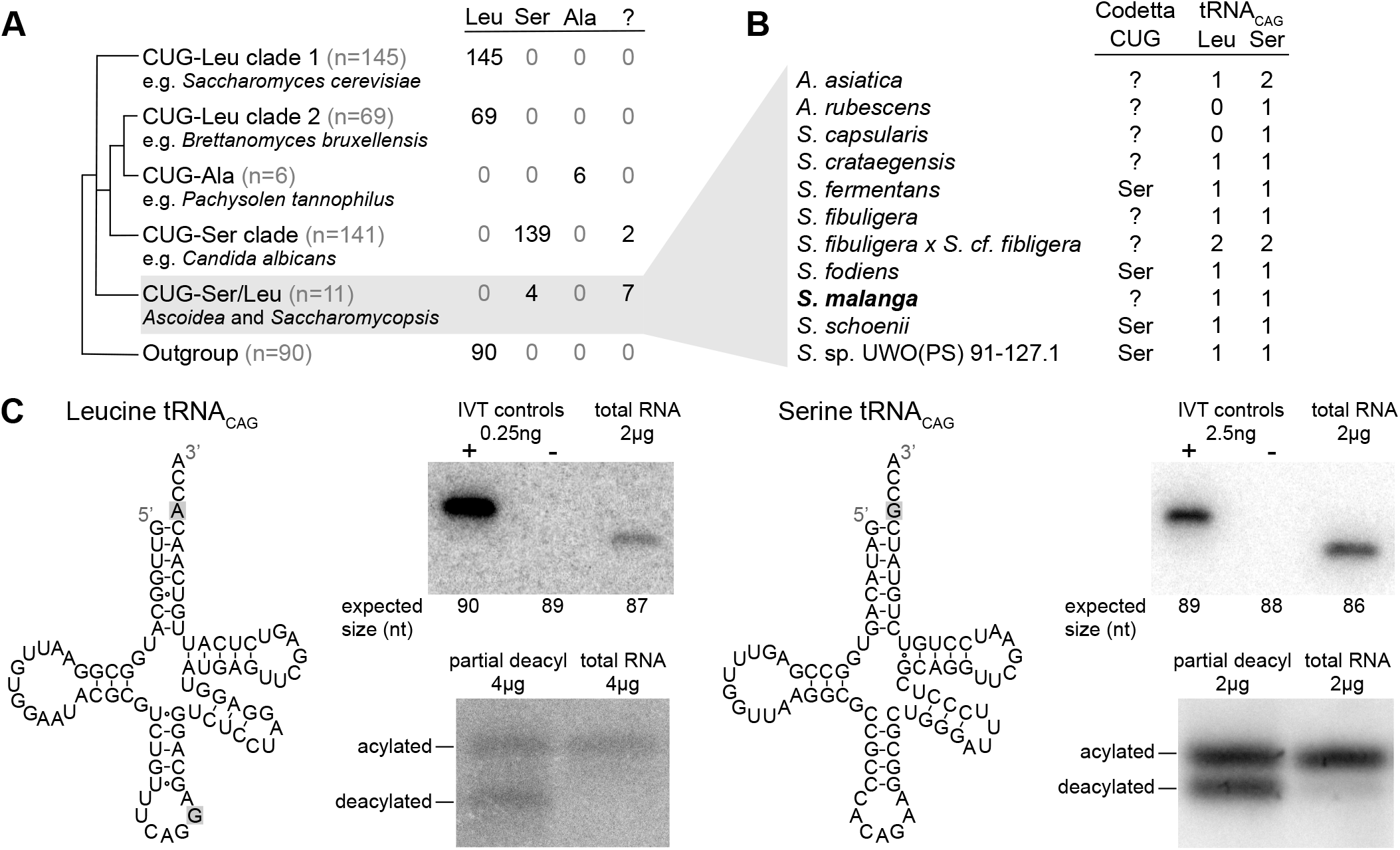
(A) CUG translation inferred by Codetta of 462 Saccharomycetes species, grouped by phylogenetic clade. Cladogram was adapted from ***Shen et al.*** (***2018***). Phylogenetic placement of CUG-Leu/Ser clade is unresolved, thus the three-way branch. (B) Codetta CUG inference and number of tRNA_CAG_ genes in *Ascoidea* and *Saccharomycopsis* genomes. tRNA_CAG_ genes were identified using tRNAscan-SE 2.0 and were classified as being serine-type or leucine-type based on the presence of tRNA identity elements. (C) Northern blotting to confirm expression and charging of leucine and serine tRNA_CAG_ genes in *S. malanga*. Probable secondary structures of the two *S. malanga* tRNA_CAG_ are shown with features used for leucine/serine classification highlighted in gray. In the tRNA expression blots, *in vitro* transcribed (IVT) versions of the target tRNA (+ control) and the most similar other tRNA (-control, as determined by sequence homology with the probe) were used as controls for probe specificity. In the tRNA charging blots, a partial deacylation control was used to help visualize the expected band sizes for acylated and deacylated versions of the probed tRNA. **Figure 2–Figure supplement 1.** Distribution of Pfam consensus column amino acid support for CUG codons in *B. inositovora* and *C. fragrans*. **Figure 2–source data 1.** Table of all analyzed yeast genomes with phylogenetic grouping and Codetta CUG inference.

The other seven species without an inferred amino acid for CUG all belong to the closely-related genera *Ascoidea* and *Saccharomycopsis* (four additional species in these clades were predicted to translate CUG as serine). Analysis of tRNA genes revealed that 9 out of 11 species in this clade encode two types of tRNA_CAG_ genes, one predicted to be serine-type and one leucine-type, suggesting that CUG may be ambiguously translated as both serine and leucine via competing tRNAs in some of these species (***Figure 2B***). We used Northern blotting to assay the expression of both tRNA_CAG_ genes in some of these species under a variety of growth conditions (data not shown), but could detect reliable expression of both serine- and leucine-type tRNA_CAG_ genes only in *Saccharomycopsis malanga* (only the serine tRNA_CAG_ could be detected in other species) (***Figure 2C***). To determine whether both tRNAs are aminoacylated, we performed acid urea PAGE Northern blotting which separates aminoacylated and deacylated tRNAs. We found that both serine and leucine *S. malanga* tRNA_CAG_ are predominantly charged in cells (***Figure 2C***), likely partaking in the translation of CUG codons. If CUG is indeed translated ambiguously in this clade, it would explain why Codetta did not place a high probability on any single amino acid decoding for some species.

The existence of serine and leucine tRNA_CAG_ genes in some *Ascoidea* and *Saccharomycopsis* yeasts was reported by ***Krassowski et al.*** (***2018***) and ***Mühlhausen et al.*** (***2018***) while we were conducting experiments. Ambiguous translation of CUG was demonstrated in *A. asiatica* (***Mühlhausen et al., 2018***); however, for *S. malanga* only expression of the serine tRNA_CAG_ could be detected (***Krassowski et al., 2018***) and incorporation of predominantly serine at protein positions encoded by CUG (***Mühlhausen et al., 2018***). In contrast to these studies, we used a saturated growth condition where the leucine tRNA_CAG_ seems to be more highly expressed. While we did not quantify the relative expression of the two tRNA_CAG_ in *S. malanga*, a visual comparison of the band intensities in ***Figure 2C*** suggests that the expression of the leucine tRNA_CAG_ is at least ten times less than the serine tRNA_CAG_ even in the saturated growth condition.

These results show that Codetta can correctly infer canonical and non-canonical codon translations and can flag unusual situations such as ambiguous translation even though it assumes unambiguous translation. All of the remaining 63 codons were inferred to use the expected translation in all species, with the following exceptions. In three species belonging to a lineage of *Hanseniaspora* with low genomic GC content (***Steenwyk et al., 2019***), the arginine codons CGC and/or CGG had a ‘?’ inference due to few (<20) aligned Pfam consensus columns. In eight other species, either the stop codon UAG or UGA was inferred to code for tryptophan due to some (<23) aligned Pfam consensus columns. We could not find any nuclear suppressor tRNA genes, and we believe these inferences are due to the erroneous alignment of Pfam domains to in-frame stop codons in pseudogenes. In-frame stop codons do not appear randomly within pseudogenes but instead are most likely to result from single nucleotide transversions from certain codons (such as the UGG tryptophan codon).

### Computational screen of all bacterial and archaeal genomes finds previously known alternative genetic codes

To explore the diversity of genetic codes in bacterial and archaeal genomes, we used Codetta to analyze 251,571 assembled genomes from GenBank, including partial assemblies and those derived from single-cell genomics and metagenomic assembly. Summaries of our analysis (***Table 1*** and ***Table 2***) are shown for a subset of the results, dereplicated to reduce the over-representation of frequently sequenced organisms by selecting a single assembly for each species-level NCBI taxonomic ID (48,693 unique species: 46,384 bacteria, 2,309 archaea). Results for the full dataset and the dereplicated subset are available in ***Table 2***-source data 1.

**Table 1.** A summary of all bacterial clades previously known to use a codon reassignment. For each clade, the NCBI taxonomic IDs (taxids) shown most closely correspond to the known phylogenetic distribution from the literature. For each codon reassignment, we show the number of sequenced species analyzed by Codetta and how many were inferred to use the expected amino acid or had no inferred amino acid. None of the analyzed species belonging to reassigned clades were predicted to use an unexpected amino acid at the reassigned codon. [1] ***Bové*** (***1993***), [2] ***Volokhov et al.*** (***2007***), [3] ***McCutcheon et al.*** (***2009***), [4] ***Bennett and Moran*** (***2013***), [5] ***McCutcheon and Moran*** (***2010***), [6] ***Salem et al.*** (***2017***), [7] ***Rinke et al.*** (***2013***), [8] ***Campbell et al.*** (***2013***)

| Phylogenetic distribution | NCBI taxids | Ref | N species | Codon reassignment | Reassigned codon |  |
| --- | --- | --- | --- | --- | --- | --- |
|  |  |  |  |  | Expected amino acid | Uninferred (?) |
| Entomoplasmatales & Mycoplasmatales | 186328, 264638, 2085 | [1, 2] | 199 | UGA Stop→W | 191 | 8 |
| *Hodgkinia cicadicola* | 573658 | [3] | 1 | UGA Stop→W | 1 | 0 |
| *Nasuia deltocephalinicola* | 1160784 | [4] | 1 | UGA Stop→W | 1 | 0 |
| *Zinderia insecticola* | 884215 | [5] | 1 | UGA Stop→W | 1 | 0 |
| *Stammera capleta* | 2608262 | [6] | 1 | UGA Stop→W | 1 | 0 |
| Gracilibacteria | 363464 | [7] | 15 | UGA Stop→G | 13 | 2 |
| Absconditabacteria | 221235 | [8] | 6 | UGA Stop→G | 6 | 0 |

**Table 2.** A summary of codon inferences from the set of genomes analyzed by Codetta, dereplicated to one assembly per species. The Codetta inference for each codon is compared against a genetic code annotation derived by layering the known bacterial genetic codes in ***Table 1*** over the NCBI taxonomy. Reassigned stop codons are included with sense codons. **Table 2–source data 1.** Table of all analyzed genome assemblies with genetic code inferred by Codetta, inclusion in the dereplicated dataset, and number of expected, unexpected, and ‘?’ codon inferences.

|  |  | Bacteria |  | Archaea |  |
| --- | --- | --- | --- | --- | --- |
|  |  | 46,384 species |  | 2,309 species |  |
| Sense | Total (N codons x N species) | 2,829,648 |  | 140,849 |  |
|  | Expected amino acid | 2,823,497 | 99.78% | 140,631 | 99.85% |
|  | Other amino acid | 612 | 0.02% | 0 | 0.00% |
|  | Uninferred (?) | 5,539 | 0.20% | 218 | 0.15% |
| Stop | Total (N codons x N species) | 138,928 |  | 6,927 |  |
|  | Amino acid | 290 | 0.21% | 9 | 0.13% |
|  | Uninferred (?) | 138,638 | 99.79% | 6,918 | 99.87% |

To see if our screen recovered known alternative genetic codes, we collated a comprehensive literature summary of all bacterial and archaeal clades known to use alternative genetic codes (***Table 1***) and layered it over the NCBI taxonomy, annotating all remaining organisms with the standard genetic code. This resulted in a genetic code annotation for each species. For most species using known alternative genetic codes in our dataset, our predictions at the reassigned codon agreed with the the expected amino acid translation (***Table 1***). There were no instances of reassigned codons predicted to translate as an unexpected amino acid, but there were a few cases of reassigned UGA codons which had no amino acid meaning inferred (‘?’ inference).

Since the uninferred codons could represent a lack of sensitivity by Codetta, we looked more closely at these examples. In the Mycoplasmatales and Entomoplasmatales, which are believed to translate the canonical stop codon UGA as tryptophan, eight species had no inferred amino acid meaning for UGA due to fewer than 4 aligned Pfam consensus columns. All of these genomes lack a UGA-decoding 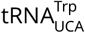 gene and all but one instead contain a release factor 2 gene (which terminates translation at UGA). Five of these species are included in the Genome Taxonomy Database (GTDB) (***Parks et al., 2020***), a comprehensive phylogeny of over 190,000 bacterial and archaeal genomes, where they are grouped into a different order (GTDB order RF39). We therefore attribute at least 5 (and perhaps all 8) as a taxonomic misannotation in the NCBI database, and we believe that UGA is a stop codon in these species. In the Gracilibacteria, which are believed to translate the stop codon UGA as glycine, two species had no inferred amino acid meaning for UGA. Neither genome contained the expected UGA-decoding 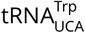 gene and both instead encoded a release factor 2 gene, supporting that UGA is a stop codon and not a glycine codon in these species. Indeed, one of these species is included in the GTDB and is grouped in a different order than the other UGA-reassigned Gracilibacteria and Absconditabacteria.

Across the 48,693 genomes (dereplicated to one assembly per species), we predicted the amino acid translation of a total of 2,970,497 individual sense codons (roughly 61 times the number of genomes), with 99.79% of the predictions consistent with the expected amino acid (similar proportion across bacteria and archaea) (***Table 2***). About 0.19% of sense codons had a ‘?’ inference, demonstrating that entire genomes contain more than enough information to infer the amino acid translation of most sense codons. Unexpected amino acid meanings were predicted for 612 sense codons. These are candidates for new codon reassignments, but could also include inference errors. For stop codons, 99.80% out of total of 145,855 stop codons across the dereplicated bacterial and archaeal genomes had no inferred amino acid meaning, as expected. 290 bacterial stop codons and 9 archaeal stop codons were inferred to translate as an amino acid, adding to our list of candidate new genetic codes.

### Validation of candidate new alternative genetic codes

To prioritize high-confidence novel genetic codes, we gathered additional evidence by examining 1) the translational components (tRNA and/or release factor genes) involved in the reassignment, 2) the usage of the reassigned codon, including manual examination of alignments of highly conserved single-copy genes, and 3) the phylogenetic extent of the proposed reassignment. Since many candidate genetic codes were found in uncultivated clades with only rough taxonomic classification on NCBI, we explored phylogenetic relationships using the Genome Taxonomy Database (GTDB) (***Parks et al., 2020***). The GTDB is a phylogeny of over 190,000 archaeal and bacterial genomes, providing provisional domain-to-species phylogenetic classifications for uncultivated as well as established clades. A list of all candidate novel genetic codes can be found in Supplementary file 1. We focused on the candidate codon reassignments with the highest degree of additional evidence and attempted to characterize common sources of error. The set of lower-confidence candidates may still include additional real codon reassignments requiring further validation.

The most common error was the inference of AGA and/or AGG arginine codons as coding for lysine, occurring in 567 bacterial species. Almost all of the AGA- and AGG-decoding tRNAs found in these genomes were consistent with arginine identity (based on the arginine identity elements A/G73 and A20), supporting that AGA and AGG are arginine codons in the majority of these species. The unusually high GC content of these genomes (ranging between 0.52 - 0.77, median 0.68) suggests that the source of the lysine inference comes from high GC content-driven nonsynonymous substitutions of the AAA and AAG lysine codons to AGA and AGG arginine codons at protein residues that can tolerate either positively-charged amino acid. As a result, AGA and AGG codons consistently appear at residues conserved for lysine in other species, which Codetta mistakes for the signature of codon reassignment. Bacteria with high genomic GC content have long been observed to preferentially use more arginine and less lysine in cellular proteins (***Sueoka, 1961***), most likely due to substitutions between the aforementioned lysine and arginine codons (***Singer and Hickey, 2000***; ***Knight et al., 2001b***). This error could be mitigated in future analyses by using profile HMMs built from sequences that match the analyzed genome in GC content or amino acid composition.

Some erroneous stop codon inferences resulted from genome contamination by organisms with known stop codon reassignments. We suspected contamination when the Pfam consensus columns aligned to a stop codon were only present in a limited part of the genome and confirmed the origin of these regions by homology search of the genes containing the in-frame stop codons. We have found examples of predicted stop reassignments in *Sulfolobus* assemblies caused by contamination with UGA-recoding *Mycoplasma* contigs, in an alphaproteobacteria assembly caused by contamination with UAA- and UAG-recoding ciliate contigs, in *Chlorolexi* assemblies caused by contamination with UGA-recoding Absconditabacteria contigs, and in others.

We found five clades using candidate novel alternative genetic codes with a convincing level of additional support, including tRNA genes that would enable the new translation. All five new genetic codes involve the reassignment of arginine codons, representing the first sense codon reassignments in bacteria.

### Reassignment of the canonical arginine codon AGG to methionine in a clade of uncultivated Bacilli

Eight bacterial genomes were inferred to translate AGG, a canonical arginine codon, as methionine. All eight genomes were assembled from fecal metagenomes of baboons or humans (***Parks et al., 2017***; ***Almeida et al., 2019***) and have only coarse-grained NCBI genome classification as uncultured Bacillales or Mollicutes bacteria. The GTDB assigns these eight genomes to a three species clade within the placeholder genus UBA7642 (family CAG-288, order RFN20, class Bacilli), of which all other species were inferred to translate AGG as arginine (***Figure 3A***).

**Figure 3.**
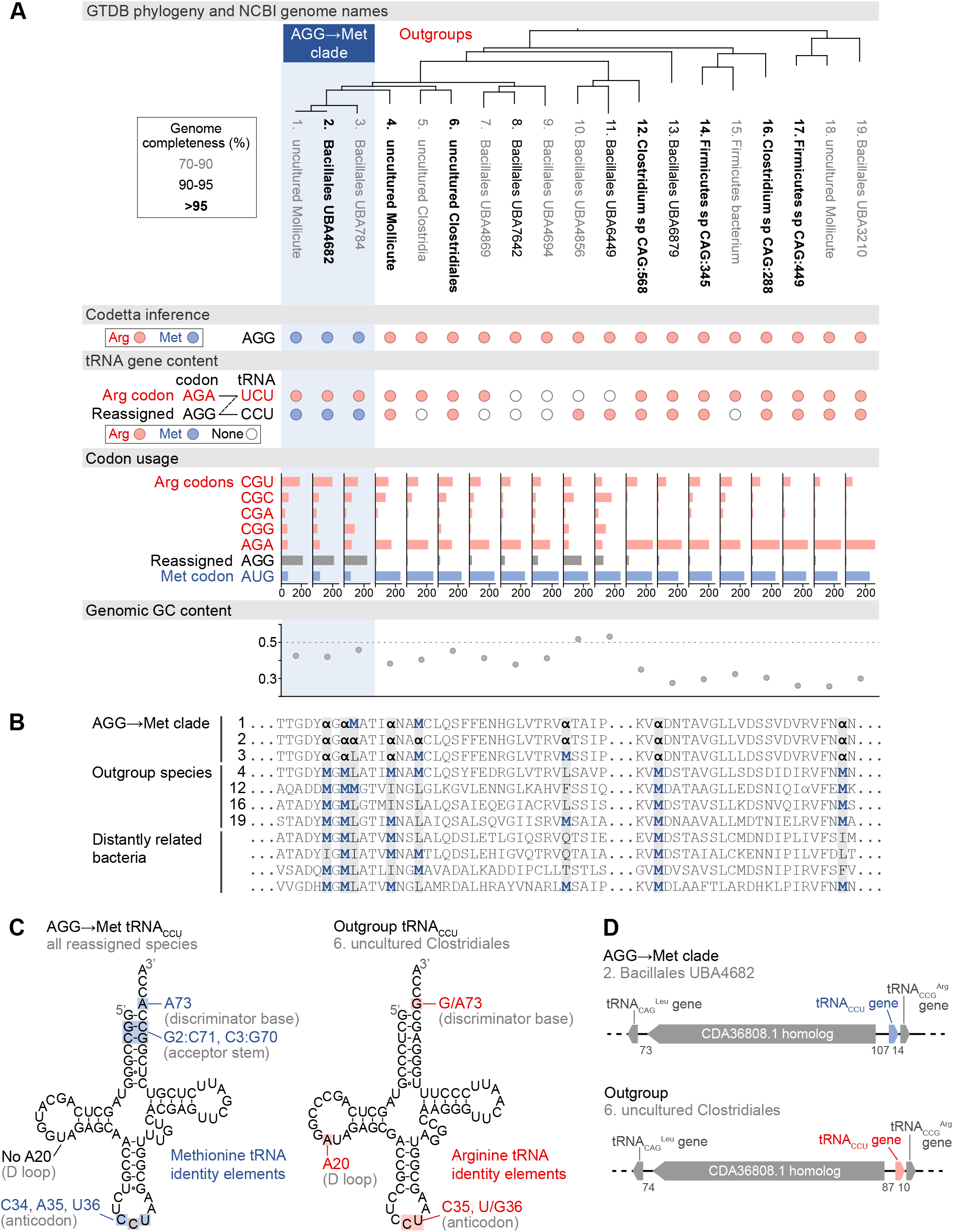
Reassignment of AGG from arginine to methionine in a clade of uncultivated Bacilli. (A) GTDB phylogenetic tree of the Bacilli AGG→Met clade and closest outgroup genomes, with the annotated NCBI genome name shaded according to the GTDB CheckM estimated genome completeness. GTDB genus UBA7642 corresponds to species #1-9 and GTDB family CAG-288 corresponds to species #1-16. For each genome, the Codetta AGG inference is indicated by colored circles (red: arginine, blue: methionine). The presence of tRNA genes is also indicated by filled circles for tRNA_UCU_ and tRNA_CCU_, colored by the predicted amino acid charging based on known identity elements (see Methods), or a white circle if no tRNA gene could be detected. The lines connecting codons and anticodons represent the likely wobble decoding capabilities, with dashed lines representing likely weaker interactions. Codon usage is the frequency per 10,000 codons aligned to Pfam domains. (B) Multiple sequence alignment of uridylate kinase (BUSCO POG091H02JZ) from the reassigned species, selected outgroup species, and four more distantly related bacteria (*Bacillus subtilis*, *Nostoc punctiforme*, *Chlamydia caviae*, and *E. coli*). All AGGs are represented by ***α***. Alignment regions containing multiple nearby AGG positions in the reassigned species are shown. (C) A comparison of the AGG-decoding tRNA_CCU_ in the Bacilli AGG → Met clade (identical sequence in all genomes) and in an outgroup genome (#6, uncultured Clostridiales). tRNA sequence features involved in methionine identity in the reassigned clade tRNA_CCU_ and arginine identity in the outgroup tRNA are highlighted (***Meinnel et al., 1993***; ***Giegé et al., 1998***), with nucleotide numbering following the convention of ***Sprinzl et al.*** (***1998***). The C35 anticodon nucleotide in the AGG→Met clade tRNA is highlighted in gray because it does not match the A35 methionine identity element. (D) The genomic context surrounding the tRNA_CCU_ gene in a member of the Bacilli AGG→Met clade (#2, Bacillales UBA4682) and in an outgroup species (#6, uncultured Clostridiales). Gene lengths and intergenic distances are drawn proportionally, with the number of base pairs between each gene indicated below. **Figure 3–source data 1.** Table of genome accessions, Codetta AGG inference, tRNA gene presence, codon usage, and genome GC content for the reassigned AGG→Met Bacilli and outgroup species shown on tree.

In each of the reassigned genomes, the AGG inference by Codetta is based on a sufficiently large number of aligned Pfam consensus columns (over 2,200 compared to an average of about 1,800 for each of the other 60 sense codons) from over 480 different Pfam domains. ***Figure 3B*** shows an example multiple sequence alignment of uridylate kinase, a single-copy conserved bacterial gene, from the reassigned species, outgroup genomes, and several more distantly related bacteria. In the reassigned clade, AGG codons are used interchangeably with AUG methionine codons and tend to occur at positions conserved for methionine and other nonpolar amino acids in the other species.

In the reassigned clade, AGG is the dominant methionine codon with a usage of 209-235 per 10,000 codons in Pfam alignments, outnumbering the canonical methionine codon AUG (59-69 per 10,000 codons) (***Figure 3A***). The process of codon reassignment involves genome-wide codon substitutions to remove the reassigned codon from positions that cannot tolerate the new amino acid, leading to depressed codon usage. High usage of AGG in the reassigned clade suggests that this is an established codon reassignment that has had time to rebound in frequency through synonymous substitutions with the standard AUG methionine codon. In many outgroup genomes, AGG is a rare arginine codon (***Figure 3A***).

Escape from viral infection has been put forth as a potential selective pressure for the evolution of alternative genetic codes, although viruses are also known to infect some alternative genetic code organisms such as *Mycoplasma* and mitochondria (***Shackelton and Holmes, 2008***). We inferred the genetic code of phage genomes assembled by ***Al-Shayeb et al.*** (***2020***) from the same baboon fecal metagenomic dataset as some reassigned Bacilli genomes. Two phage assemblies were predicted to translate AGG as methionine (assemblies GCA_902730795.1 and GCA_902730815.1). The assemblies do not contain genes for the AGG-decoding tRNA_CCU_, so the phage presumably rely on the host tRNAs for translation. Thus, some phage may have adapted to the AGG translation as methionine in the reassigned Bacilli.

We used tRNAscan-SE 2.0 (***Chan et al., 2019***) to determine which tRNAs are available to decode AGG in the reassigned and outgroup genomes (***Figure 3A***). Some tRNA genes are missing, possibly due to the incomplete nature of some metagenome-assembled genomes as indicated by low genome completeness estimates. The cognate tRNA for the AGG codon, tRNA_CCU_, from the reassigned clade has features of methionine identity (including an A73 discriminator base and G2:C71 and C3:G70 base pairs in the acceptor stem) and lacks the important arginine identity element A20 in the D-loop (***Meinnel et al., 1993***; ***Giegé et al., 1998***), supporting translation of AGG as methionine (***Figure 3C***). *In vitro* experiments have shown that anticodon mutations to 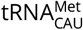 disrupt recognition by the methionyl-tRNA synthetase in *E. coli*; however, the C35 change necessary to decode the AGG codon affects the least critical anticodon nucleotide (***Schulman and Pelka, 1983***). The outgroup genomes contain a tRNA_CCU_ with features of arginine identity (including a G73 discriminator base and A20 in the D-loop). The genomic context of the tRNA_CCU_ is similar in many reassigned clade and outgroup genomes, flanked by a 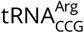 immediately downstream and a homolog of GenBank protein CDA36808.1 upstream (***Figure 3D***). This implies that the reassigned and outgroup tRNA_CCU_ evolved from the same ancestral tRNA gene, and the reassigned methionine tRNA_CCU_ likely emerged through a change in aminoacylation of an arginine tRNA_CCU_ rather than through duplication and anticodon mutation of a methionine tRNA.

The reassigned genomes use an arginine-type tRNA_UCU_ to decode the unaffected AGA arginine codon. Depending on the post-transcriptional modification of the U34 anticodon nucleotide, the arginine tRNA_UCU_ could recognize AGG via wobble and potentially cause ambiguous translation. In *E. coli*, the U34 of tRNA_UCU_ is modified to 5-methylaminomethyluridine (***Sakamoto et al., 1993***) which primarily decodes the AGA codon with a low level of AGG recognition (***Spanjaard et al., 1990***). ***Mukai et al.***(***2015***) demonstrated that it is possible to engineer separate decodings for AGA and AGG in *E. coli* by reducing expression level of the tRNA_UCU_ to the point where decoding of AGG by tRNA_UCU_ is presumably insignificant in competition with the cognate tRNA_CCU_. In most outgroup genomes, AGA is the dominant arginine codon, while in the reassigned clade the preferred arginine codon is CGU (***Figure 3A***), which may indicate reduced demand and expression of tRNA_UCU_ to avoid ambiguous translation of AGG. A similar potential for ambiguous translation due to U34 wobble exists with the previously known decoding of UGA as glycine and UGG as tryptophan in Absconditabacteria and Gracilibacteria (***Campbell et al., 2013***; ***Rinke et al., 2013***).

### Reassignments of arginine codons CGA and CGG occur in clades with low genomic GC content

The remaining four clades with high-confidence codon reassignments all affect the arginine codons CGA and/or CGG (***Figure 4***). Three clades are in the phylum Firmicutes: the genus *Peptacetobacter* is predicted to translate CGG as glutamine (***Figure 4–Figure Supplement 1***), a clade of uncultivated Bacilli in the GTDB order RFN20 (same as the AGG-reassigned Bacilli) is predicted to translate CGG as tryptophan (***Figure 4–Figure Supplement 2***), and members of the genus *Anaerococcus* are also predicted to translate CGG as tryptophan (***Figure 4–Figure Supplement 3***). The fourth clade is Absconditabacteria (also known as Candidate Division SR1, part of the Candidate Phyla Radiation), which is predicted to have reassigned CGA and CGG both to tryptophan (***Figure 4–Figure Supplement 4***), in addition to the already known reassignment of UGA from stop to glycine.

**Figure 4.**
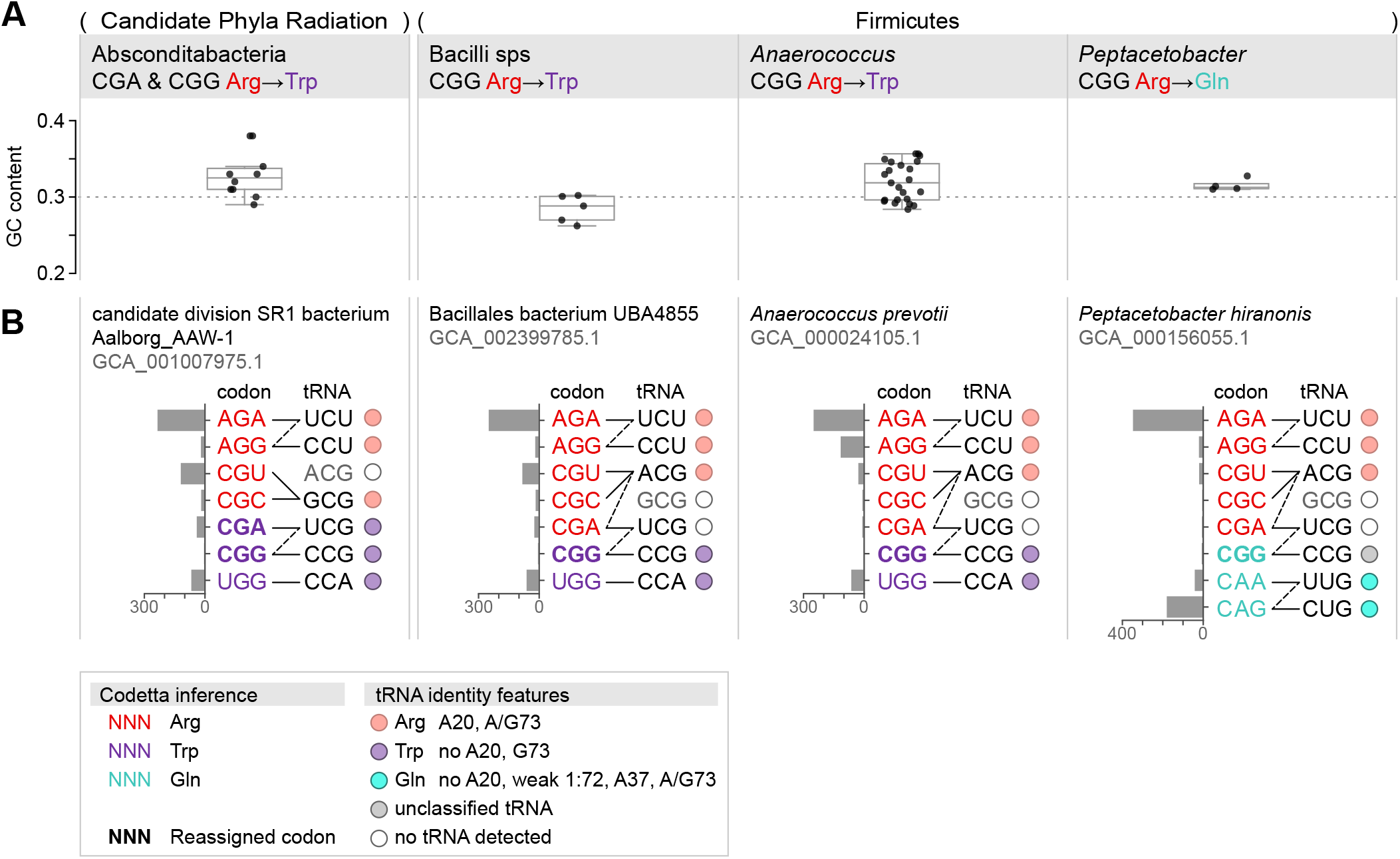
Summary of GC content, codon usage, and tRNA genes of four CGA and/or CGG reassignments. (A) Distribution of genomic GC contents across all species in the reassigned clades. (B) For each reassigned clade, we selected a representative species to show codon usage and tRNA decoding ability. Codon usage is plotted for the reassigned codon and for all other codons of the original and new amino acids in usage per 10,000 in Pfam alignments. Codons are colored by their Codetta inference and reassigned codons are bolded. The lines connecting codons and tRNA anticodons represent the likely wobble decoding capabilities, with dashed lines representing likely weaker interactions. The anticodon ACG is presumed to be modified to ICG, and UCG is presumed to be modified in a way that restricts wobble to CGA and CGG, but could potentially recognize CGU and CGC as well depending on the true modification state. Anticodons in gray font are not expected to be found in the respective clade. Presence of tRNA genes is indicated by filled circles, colored by the predicted amino acid charging based on the identity elements in the key. **Figure 4–Figure supplement 1.** Reassignment of CGG→Gln in *Peptacetobacter*. **Figure 4–Figure supplement 2.** Reassignment of CGG→Trp in a clade of Bacilli. **Figure 4–Figure supplement 3.**Reassignment of CGG→Trp in *Anaerococcus*. **Figure 4–Figure supplement 4.** Reassignment of CGA and CGG→Trp in Absconditabacteria. **Figure 4–source data 1.** Table for each reassignment containing genome accessions, Codetta CGA/CGG inference, tRNA gene presence, codon usage, and genome GC content for the reassigned clade and outgroup species.

In contrast to the reassignment of AGG to become the dominant methionine codon (described in the previous section), these CGA/CGG reassignments resemble earlier stages of codon reassignment where the reassigned codon has not yet rebounded in frequency through synonymous substitutions with the new amino acid. Due to the rarity of the reassigned CGA/CGG codons, these predictions are based on fewer aligned Pfam consensus columns and may be more prone to error. As a check for each reassignment, we looked for examples of the reassigned codon in conserved regions of single-copy gene alignments (***Figure 4–Figure Supplement 1B - Figure 4–Figure Supplement 4B***) and found multiple supporting positions for all reassigned codons except the extremely rare CGG codon in *Anaerococcus*. We also looked for tRNA genes with an anticodon and amino acid identity elements consistent with the reassignment (***Figure 4–Figure Supplement 1 - Figure 4– Figure Supplement 4***), and found consistent tRNAs for all clades except for *Peptacetobacter* whose CGG-decoding tRNA_CCG_ resembles neither an arginine or glutamine isotype. While amino acid conservation at the reassigned codon and sequence-based prediction of tRNA charging may lend support to a predicted codon reassignment, only experimental confirmation can establish how the reassigned codons are translated *in vivo* and whether there is ambiguous translation. In particular, *Anaerococcus* and *Peptacetobacter* include culturable species and may be experimentally confirmed in the future.

The four CGA/CGG candidate reassignments share several features that suggest common evolutionary forces at play. Most notable is the very low genomic GC content of the reassigned clades (0.26-0.38, ***Figure 4A***). In all four clades, the usage of GC-rich CGN-box codons–including CGA and CGG–is depressed and arginine residues are primarily encoded by AGA codons (***Figure 4B***). In the three Firmicute CGG reassignments, CGG is an extremely rare codon (codon usage <6 per 10,000 in aligned Pfam domains for all species). In the Absconditabacteria, CGG also tends to be quite rare (<7 per 10,000 in all but one species) with CGA slightly more abundant (<37 per 10,000 in all species). In one Absconditabacteria (assembly GCA_002791215.1), the frequency of both CGA and CGG approaches the frequency of the canonical tryptophan codon UGG, consistent with a more advanced stage of codon reassignment (usage of CGA and CGG is 30 and 24 per 10,000, compared to 35 for UGG). Low genomic GC content is thought to be created by mutational bias in favor of AT nucleotides, causing a gradual shift towards synonymous codons with lower GC compositions (***Knight et al., 2001b***; ***Muto and Osawa, 1987***). This may have helped disfavor usage of CGA and/or CGG prior to reassignment, lessening the impact of changing the codon meaning.

The tRNAs used to decode the CGN codon box may have also influenced the reassignment of CGA and CGG codons. A shared feature of the three Firmicute CGG reassignments is that the tRNA_UCG_ is missing (***Figure 4B***), presumably lost prior to or during the reassignment of CGG. If the tRNA_UCG_ were present, it would likely recognize both CGA and CGG via wobble which would complicate assigning different amino acid meanings to those two codons. In the absence of tRNA_UCG_, CGA (along with CGU and CGC) is presumably decoded by a 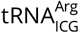 (derived by deamination of 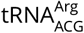 ACG, the only widespread instance of inosine tRNA wobble in bacteria). This leaves CGG to be decoded solely by a tRNA_CCG_ (***Figure 4B***). In this situation, CGG is one of a few codons in the genetic code decoded by a single dedicated tRNA, potentially facilitating codon reassignment since the translational meaning of CGG can now be altered independently of neighboring codons. The inosine wobble modification is not used by some deeply branching bacteria (***Rafels-Ybern et al., 2018***), and the 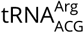 gene appears to be lacking in the Candidate Phyla Radiation, including Absconditabacteria. Instead, these bacteria use a 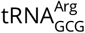 to decode CGU and CGC, and rely on a tRNA_UCG_ and tRNA_CCG_ to recognize CGA and CGG (***Figure 4B***). Since the ability of tRNA_UCG_ to decode CGA and CGG makes it difficult to split the translational meanings of the two codons, it may explain why both CGA and CGG are reassigned to tryptophan together in the Absconditabacteria.

For some of these reassignments, close outgroup species may shed light on potential intermediate stages of codon reassignment. The CGG reassignment in the Absconditabacteria may extend to members of the sister clade Gracilibacteria– some Gracilibacteria were predicted to translate CGG as tryptophan, while others translate CGG as arginine (***Figure 4–Figure Supplement 4***). This may reflect a complicated history of CGG reassignment and possible reversion to arginine translation. For the CGG reassignment in *Peptacetobacter*, the closest sister group (which includes the pathogen *Clostridioides difficile*) has extremely rare usage of CGG (<1 per 10,000 in aligned Pfam domains in all but two species) and appears to lack any tRNA capable of decoding CGG by standard codon-anticodon pairing rules (***Figure 4–Figure Supplement 1***). This may resemble an intermediate stage in codon reassignment before the ability to translate CGG as a new amino acid is gained, similar to the unassigned CGG codon in *Mycoplasma capricolum* (***Oba et al., 1991***). In *Anaerococcus*, all species contain a CGG-decoding tRNA_CCG_ with features of tryptophan identity (***Figure 4–Figure Supplement 3***). Unexpectedly, members of an outgroup genus *Finegoldia* also have a tRNA_CCG_ with features of tryptophan identity (CGG inferred to be ‘?’ by Codetta). It is unclear if the tRNA_CCG_ genes in these two clades share an evolutionary history or represent independent events.

## Discussion

We present a method for computationally inferring the genetic code that can scale to analyze hundreds of thousands of genomes which we call Codetta. We validate Codetta on the well-studied reassignments of CUG in yeasts and rediscover the ambiguous translation of CUG as serine and leucine in *Ascoidea* and *Saccharomycopsis* by two differently charged tRNAs. We conduct the first systematic survey of genetic code usage across the majority of sequenced organisms, analyzing all sequenced bacteria and archaea (over 250,000 assemblies). The five new alternative genetic codes described here substantially expand the known diversity of codon reassignments in bacteria. Now, in addition to reassignments of the stop codon UGA to tryptophan or glycine, we have the first sense codon reassignments in bacteria, affecting the arginine codons AGG, CGA, and CGG. Two reassignments occur in culturable bacteria– in *Anaerococcus* and *Peptacetobacter*–and could be experimentally confirmed in the future, for example by proteomic mass spectrometry.

Since Codetta selects the most likely amino acid translation among the twenty canonical amino acids, some types of codon reassignments may be missed. We cannot predict reassignment to a noncanonical amino acid– for such codons, Codetta would pick the non-specific model or an amino acid that is used similarly in other species. We also cannot directly detect ambiguous translation, which may represent an important stage in codon reassignment. However, the failure to infer an amino acid translation despite a significant number of aligned Pfam consensus columns may hint at ambiguous translation, as was the case for CUG in *Ascoidea* and *Saccharomycopsis*. Since we do not model translational initiation and termination, we cannot detect the use of new start and stop codons or context-dependent stop codons that also possess an amino acid meaning, known to occur in some eukaryotes (***Swart et al., 2016***; ***Heaphy et al., 2016***; ***Záhonová et al., 2016***).

Expanding our analysis to eukaryotic, organellar, and viral genomes will help fill in the diversity of alternative genetic codes, but poses additional challenges. Since we align profile HMMs to a six-frame translation of the entire genome, the pervasive pseudogenes in many eukaryotic genomes will likely increase the rate of incorrect codon inferences by having sufficient homolgy for alignment but enough accumulated mutations to cause incorrect pairing of codons to consensus columns. Smaller scale surveys of eukaryotic genetic code diversity have focused on transcriptomic datasets (***Swart et al., 2016***; ***Heaphy et al., 2016***), which may alleviate this problem. Some viral and organellar genomes have very few protein-coding genes which may limit the ability to confidently infer the entire genetic code. One strategy is to improve sensitivity at the cost of generalizability by using clade-specific profile HMMs instead of Pfam, which may increase the proportion of aligned coding sequence. Another challenge in some organellar genomes is extensive mRNA editing (***Gray, 1996***; ***Alfonzo et al., 1997***), which violates our assumption that the genomic codon sequence represents the mRNA sequence and may require analyzing the edited transcriptome to ensure correct correspondence of codons to profile HMM positions.

In the ‘codon capture’ model of codon reassignment, genome-wide pressures such as biased GC content or genome reduction drive a codon to near extinction such that the codon can acquire a new tRNA decoding with a minimal effect on translation (***Osawa and Jukes, 1989***). Most UGA reassignments in bacteria occur in clades with very low genomic GC content, which is thought to have reduced UGA to very low usage in favor of the stop codon UAA. This includes the Mycoplasmatales and Entomoplamatales (0.24-0.39 GC) (***Jukes, 1985***; ***Bové, 1993***), Absconditabacteria and Gracilibacteria (0.21-0.53 GC, ***Figure 4***–***source data 1***) (***Campbell et al., 2013***; ***Rinke et al., 2013***), and most insect endosymbiotic bacterial reassignments (0.13-0.17 GC, with the notable exception of *Hodgkinia cicadicola* with 0.58 GC) (***McCutcheon and Moran, 2010***; ***Bennett and Moran, 2013***; ***Salem et al., 2017***; ***McCutcheon et al., 2009***). The CGA and/or CGG reassignments described here similarly exhibit low genomic GC content (0.26-0.38) and very rare usage of GC-rich codons including CGA and CGG. A codon does not need to completely disappear for reassignment to be facilitated by rare codon usage, and it is likely that a brief period of translational ambiguity or inefficiency helps drive the remaining codon substitutions. We posit that, in bacteria, reduction in codon usage driven by genome-wide processes plays a major role in enabling codon reassignment and may explain why codon reassignments repeatedly evolve in clades such as Firmicutes (known for their low genomic GC content) and lifestyles such as endosymbiosis (which is often accompanied by genome reduction and skewed GC content) (***McCutcheon and Moran, 2011***).

All five of the new reassignments affect arginine codons (AGG, CGA, and CGG). While these are the first instances of arginine codon reassignment in non-organellar genomes, several arginine reassignments are known in mitochondria: in various metazoan mitochondria the codons AGA and AGG have been reassigned to serine, glycine, and possibly stop and AGG has been reassigned to lysine (***Knight et al., 2001a***; ***Abascal et al., 2006a***), and in various green algal mitochondria AGG has been reassigned to alanine and methionine and CGG to leucine (***Noutahi et al., 2019***). Arginine codons have several unique features that may predispose them to codon reassignment. First, across the tree of life, arginine has an over-representation of codons in the genetic code relative to usage in proteins (***Jukes et al., 1975***; ***King and Jukes, 1969***), contributing to rare usage of the least favored arginine codon. Second, the six arginine codons range from one to three GC nucleotides in composition (only equaled by leucine), which may create greater bias in codon usage in response to genomic GC content than for amino acids with less GC variability in their codons. In organisms with small genomes, these features alone might make the rarest arginine codon very low in number and more susceptible than other codons to reassignment. The arginine codon CGG may be even more of a target for reassignment because, in most bacteria, the only widespread instance of inosine tRNA wobble is used to decode the CGU, CGC, and CGA arginine codons (***Grosjean et al., 2010***). In the absence of a tRNA_UCG_, CGG is decoded by a dedicated tRNA_CCG_ and can be reassigned without affecting the translation of other codons.

Some codon reassignments have convergently reappeared across the tree of life: CGG to tryptophan in three bacterial clades described here, AGG to methionine in a clade of Bacilli described here and in green algal mitochondria (***Noutahi et al., 2019***), UGA to tryptophan in multiple bacterial, mitochondrial, and eukaryotic lineages (***Knight et al., 2001a***), and others. Recurrent changes could reflect 1) a common evolutionary process, e.g. low GC content-driven reassignments disproportionately affecting codons sensitive to GC fluctuations, or 2) shared constraints imposed by conserved translational machinery, including tRNAs and aminoacyl-tRNA synthetases. For example, the tRNA anticodon-codon pairing rules dictate that U- and C-ending codons cannot be assigned separate meanings, and indeed this has not been observed in any known genetic codes. This may explain why in low GC content genomes, we see reassignments of the arginine codon CGG but not the arginine codon CGC, which would have to be reassigned together with CGU. The selection of amino acid changes in the codon reassignments described here is not clearly explained by biochemical similarity (except possibly for the reassignment of CGG from arginine to glutamine). The amino acid choice may be related to the constraints on evolving new tRNA anticodons. Most of the changes described here (and indeed all of the changes known in bacteria) involve a single nucleotide difference from cognate anticodons: tRNA_CCU_ in addition to 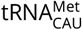 for the AGG to methionine reassignment, tRNA_CCG_ in addition to 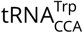 for the CGG to tryptophan reassignments, and tRNA_CCG_ in addition 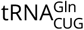 for the CGG to glutamine reassignment. Evolving a new anticodon through a single mutation may be more probable than through multiple mutations. However, the methionine tRNA_CCU_ involved in the reassignment of AGG in a clade of Bacilli appears to have evolved from an arginine tRNA_CCU_ through mutations that altered aminoacyl-tRNA synthetase recognition, rather than by an anticodon mutation to a methionine tRNA_CAU_ gene. Alternatively, this pattern could result from a limitation on the new anticodons that an aminoacyl-tRNA synthetase could accept, since most aminoacyl-tRNA synthetases use the anticodon in part to distinguish cognate and non-cognate tRNAs (***Giegé et al., 1998***). Upon characterizing the diversity of genetic codes in other parts of the tree of life, we may discover that the general patterns and evolutionary pressures differ from bacteria, reflecting differences in translational machinery, lifestyle, or genome characteristics.

## Methods

### Computational inference of the genetic code from nucleotide sequence

A preliminary translation of the input nucleotide sequences is produced by first breaking any long sequences into nonoverlapping 100 Kb pieces (because of a limit on input protein sequence length for `hmmscan`), then translating into all six frames (as six polypeptide sequences) using the standard genetic code with stop codons translated as ‘X’. A custom version of Pfam 32.0 profiles was produced from Pfam seed alignments using `hmmbuild --enone`, which turns off entropy weighting, resulting in emission probability parameters closer to the original amino acid frequencies in the input alignments. Significant homologous alignments were identified by searching each translated polypeptide against the custom Pfam database using `hmmscan` from HMMER 3.1b2 for domain hits with E-value < 10^−10^.

Alignments were further filtered to remove uncertainly aligned consensus columns (posterior probabilities of alignment <95%). By default, no single Pfam consensus columns was allowed to account for more than 1% of total aligned consensus columns for a codon, in order to mitigate some artifacts such as repetitive pseudogene families in some genomes; when this happened, the number of codon positions aligned to that specific consensus column was downsampled to 1% of the total (if a codon was aligned to fewer than 100 Pfam consensus columns total, then each unique consensus columns was downsampled to 1 occurrence). We excluded hits to five classes of Pfam models including mitochondrial proteins, viral proteins, selenoproteins, pyrrolysine-containing proteins, and proteins belonging to transposons and other mobile genetic elements. These filtered sets of aligned consensus columns defined the input 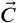 sets for each codon. The equations from the main text are then used (in log-probability calculations for numerical stability) to infer 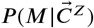 for each codon, with a default decoding probability threshold of 0.9999.

The computational requirements are dominated by the `hmmscan` step, which takes about an hour on a single CPU core for a ~12 Maa six-frame translation of a typical 6 Mb bacterial genome. We ran different genomes in parallel on a 30,000 core computing resource, the Harvard Cannon cluster. We implemented this method as Codetta v1.0, a Python 3 program that can be found at https://github.com/kshulgina/codetta/releases/tag/v1.0.

### Measuring error rate and power on synthetic datasets

A six-frame translation of the *E. coli* O157:H7 str. Sakai genome (GCA_000008865.2) was searched against the custom `--enone` Pfam 32.0 profile database as described above. We generated 400 different random subsamples each of 2, 5, 10, 20, 50, or 500 aligned consensus columns per sense codon and inferred the most likely decoding as described above. A codon inference was considered “true” (T) if the correct amino acid meaning was inferred, “false” (F) if an incorrect amino acid meaning was inferred, and “uninferred” (U) if the non-specific decoding was most probable or if no model surpassed the model probability threshold. For a given model probability threshold, per-codon error rate is the fraction of samples with a false inference (F / (T + F + U)). Per-codon power is the fraction of samples with a true inference (T / (T + F + U)). Both values were evaluated individually for each sense codon and also aggregated across all sense codons.

### Genetic code inference of archaeal and bacterial genomes

Assembly identifiers for all archaeal and bacterial genomes were downloaded from the NCBI Genome database on June 4th, 2020, and Codetta analysis was performed on all archaeal and bacterial genome assemblies. Genetic code inference results for all analyzed genomes can be found in ***Table 2***-source data 1. A variety of additional files supporting new genetic codes are available at https://github.com/kshulgina/ShulginaEddy_21_genetic_codes.

We used the NCBI taxonomy database (downloaded on July 15th, 2020) to cross-reference all assemblies with taxonomic identifiers. All analyzed genome assemblies from GenBank are associated with a NCBI taxonomic ID (taxid). Because some of these taxids correspond to subspecies or strain-level designations, we assigned a species-level taxid to each assembly by iteratively stepping up the NCBI taxonomy until a species-level node was reached. To create a dereplicated dataset, we picked one genome assembly per NCBI species-level taxid. If multiple genome assemblies were associated with an NCBI species-level taxid, assemblies were sorted based on RefSeq category (reference, representative, or neither) and then genome completeness level and a single genome assembly was randomly selected from the highest ranked category.

Phage assemblies derived from the same metagenomic samples as the AGG-recoding Bacilli were obtained by identifying the phage assemblies from ***Al-Shayeb et al.*** (***2020***) whose sample accessions were linked to the metagenomic sequencing experiments SRX834619, SRX834622, SRX834629, SRX834636, SRX834653, SRX834655, or SRX834666. Codetta analysis of the phage genomes was performed as described above.

### Cross-referencing the NCBI taxonomy with known distributions of genetic code usage

A complete list of bacterial clades previously known to use alternative genetic codes was collated with corresponding references for genetic code discovery and taxonomic distribution (***Table 1***). For each clade, we determined a set of NCBI taxids best defining the phylogenetic extent of each reassigned clade. We used this to generate a curated genetic code annotation for all NCBI species-level taxids: for the taxids defining each reassigned clade, all species-level child nodes were annotated with the alternative genetic code; all remaining species-level taxids were annotated with the standard genetic code.

We used the Genome Taxonomy Database (GTDB, version R05-RS95) (***Parks et al., 2020***) to determine the phylogenetic placement of species that use candidate new genetic codes and to identify the most closely related outgroup species.

### Identification of tRNA genes and other translational components

The tRNA gene content of genomes was determined by running tRNAscan-SE 2.0 (***Chan et al., 2019***) with default settings and a tRNA model appropriate for the domain of life (i.e. option `-E` for eukaryotes, `-B` for bacteria, `-A` for archaea). To help ensure that tRNAs of interest were not missed, we also ran a low-stringency search with the general tRNA model and no cutoff score (options `-G -X 0`) and manually examined the output.

We searched bacterial genomes for release factor genes with `hmmscan` for the TIGRFAM 15.0 (***Haft et al., 2013***) release factor 2 model (TIGR00020) and release factor 1 model (TIGR00019) against a six-frame translation of the entire genome with default settings. Since these genes are homologs, if the two models hit overlapping genomic coordinates, we kept the hit with the more significant E-value.

For the AGG arginine to methionine reassignment in a clade of Bacilli, we classified tRNA_CCU_ genes as being primarily arginine acceptors if the tRNA had A20 in the D-loop and a A/G73 discriminator base, and primarily methionine acceptors if the tRNA had an A73 disciminator base and not A20 in the D-loop (***Giegé et al., 1998***). The weaker methionine identity elements G2:C70, C3:G69 in the acceptor stem were used to support the assignment (***Meinnel et al., 1993***). In the reassignment of CGG to glutamine in *Peptacetobacter*, we classified tRNAs as arginine-type using the rules above, and as glutamine-type if the tRNA was missing arginine identity element A20 and contained the set of glutamine identity elements consisting of a weak 1:72 basepair, A37, and A/G73 (***Jahn et al., 1991***). We took the additional glutamine identity elements G2:C71 and G3:C70 in the acceptor stem, G38 in the anticodon loop, and G10 in the D-stem as support for glutamine identity (***Jahn et al., 1991***; ***Hayase et al., 1992***). For the reassignments of CGA and/or CGG arginine to tryptophan, we classified tRNAs as primarily arginine acceptors using the rules above, and provisionally as tryptophan acceptors if the tRNA had a G73 discriminator base and not A20 in the D-loop (***Giegé et al., 1998***). We considered the weak tryptophan identity element A/G1:U72 in the acceptor stem as support for tryptophan identity but did not require it (***Himeno et al., 1991***). In the Absconditabacteria and Gracilibacteria, we classified tRNA_UCA_ genes as glycine acceptors if the tRNA had G1:C72, C2:G71, G3:C70 in the acceptor stem and U73 discriminator base (***Giegé et al., 1998***). We refrained from assigning identity if the tRNA did not fit the above patterns or if the D-loop sequence was unusual such that it was unclear which nucleotide is N20. D-loop and variable loop insertions were placed at positions following the convention of ***Sprinzl et al.*** (***1998***).

### Multiple sequence alignment of BUSCO genes

For some candidate novel alternative genetic codes, we constructed multiple sequence alignments of conserved single-copy bacterial genes from the BUSCO database v3 (***Waterhouse et al., 2018***). To identify orthologs of a BUSCO gene in a particular genome, we first created a dataset of putative protein sequences by translating all open reading frames longer than 50 codons using the inferred genetic code (assuming standard stop codons unless reassigned), with candidate reassigned codons translated as ‘X’. Then, we queried each of the 148 bacterial BUSCO profile HMMs against all putative proteins using `hmmsearch` from HMMER 3.1b2 with default settings and an E-value cutoff of 10^−13^, and picked the most significant hit if it also yielded a reciprocal best hit against the entire BUSCO profile HMM database using `hmmscan` with the same E-value cutoff. Multiple sequence alignments were generated using MAFFT v7.429 (***Katoh and Standley, 2013***) with default settings.

For the described novel genetic codes, BUSCO alignments containing the reassigned codon in the reassigned clade were individually inspected and alignments containing the reassigned codon at conserved positions in well-aligned regions were preferentially selected as example alignments.

### Annotation of genomic context

To determine the genomic context surrounding the tRNA_CCU_ gene in the uncultivated Bacilli predicted to have reassigned AGG to methionine and in close outgroup genomes, we predicted tRNA and protein coding genes in the whole genome as described above. We annotated each putative protein coding gene with the reciprocal best hit homolog among annotated protein-coding genes in the outgroup assembly GCA_000434395.1 using `phmmer` from HMMER 3.1b2 with a 10^−10^ E-value cutoff.

### Phylogenetic grouping and Codetta analysis of CUG usage by budding yeasts

For analysis of CUG translation in budding yeasts, we selected all genomes belonging to the class Saccharomycetes (NCBI taxid 4891), which represent 463 unique NCBI species taxids with at least one genome. The genomes were dereplicated to one assembly per species-level taxid as described above. Yeast species were split into six taxonomic categories based on the “major clade” annotation from the phylogenetic analysis by ***Shen et al.*** (***2018***) as follows: Outgroups (major clades: Lipomycetaceae, Trigonopsidaceae, Dipodascaceae/Trichomonascaceae, Alloascoideaceae, Sporopachydermia), CUG-Leu clade 1 (major clades: Phaffomycetaceae, Saccharomycodaceae, Saccharomycetaceae), CUG-Leu clade 2 (major clade: Pichiaceae), CUG-Ser (major clade: CUG-Ser1), CUG-Ala (major clade: CUG-Ala), and CUG-Ser/Leu (major clade: CUG-Ser2). Species that were not included in the analysis by ***Shen et al.*** (***2018***) were sorted into the same major clade as other members of their annotated genus on NCBI. A single species (*Candida* sp. JCM 15000) could not be placed into a category and was excluded from the analysis. The expected CUG translation for each clade follows ***Shen et al.*** (***2018***) and is consistent with other studies of CUG translation (***Riley et al., 2016***; ***Krassowski et al., 2018***; ***Mühlhausen et al., 2018***). Genetic codes were predicted by Codetta as described above. A table describing all yeast genomes analyzed can be found in ***Figure 2***–***source data 1***.

### Identification of tRNA genes and isotype classification in yeasts

tRNA gene content of yeast genomes was determined using tRNAscan-SE 2.0 as described above. In eukaryotes, only leucine- and serine-tRNAs have a long (>12 nucleotide) variable loop so we used this feature to confirm the tRNA_CAG_ identity as serine or leucine. In yeast, serine tRNAs typically have a conserved G73 discriminator base but can tolerate any nucleotide (***Himeno et al., 1997***), while leucine tRNA identity is conferred by a A73 discriminator base and A35 and G37 in anticodon loop (***Soma et al., 1996***). We categorized tRNA_CAG_ genes as either serine-acceptors or leucine-acceptors based on the presence of these features. In some CUG-Ser clade species, serine CAG-tRNAs containing a G37 have been found to be charged with leucine at a low level (3%) (***Suzuki et al., 1997***); for categorization purposes, we would consider these tRNAs to be primarily serine-acceptors.

### *S. malanga* growth and RNA extraction

*S. malanga* (NRRL Y-7175) was obtained from the Agricultural Research Service Culture Collection (Peoria, Illinois USA). Cells were inoculated into 5 mL of YPD liquid media (containing 1% yeast extract, 2% peptone, and 2% dextrose) from a colony on a YPD agar plate and grown to saturation for 4 days at 25°C on rotating wheel.

Total RNA was extracted in acidic conditions to preserve tRNA charging, following the steps outlined in ***Varshney et al.*** (***1991***) with the following modifications. Cells were harvested by centrifugation (5 minutes at 4,000 rpm at 4°C), resuspended in 500 *μ*L ice cold buffer containing 0.3 M NaOAc pH 4.5 and 10 mM EDTA and added to 500 *μ*L ice cold phenol:chloroform (pH 4.5) and 500 *μ*L of 0.4-0.5*μ*m acid washed glass beads for cell lysis. All RNA extraction steps were performed at 4°C. In the first round of extraction, cells were vortexed for 30 minutes, rested on ice for 3 minutes, centrifuged for 15 minutes at 20,000×*g*, and the aqueous layer was transfered to 500 *μ*L of phenol:chloroform (pH 4.5), which was subject to a second round of extraction (identical, except for 3 minute vortex). A last round of extraction was performed in 500 *μ*L of chloroform with a 15 second vortex and 2 minute centrifugation. RNA in the aqueous phase was precipitated and resuspended in buffer containing 10 mM NaOAc pH 4.5 and 1 mM EDTA.

### Northern blotting for tRNA expression

The single-stranded DNA probes used for detection of *S. malanga* 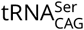 (5’ GAAATCCCAGCGC-CTTCTGTGGGCGGCGCCTTAACCAAACTCGGC 3’) and *S. malanga* 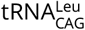 (5’ TTGACAATGAGACTC-GAACTCATACCTCCTAG 3’) were 5’ end-labelled with [*y*-P^32^]-ATP by T4 polynucleotide kinase (New England Biosciences) and purified using ProbeQuant G-50 Micro Columns (GE Healthcare Life Sciences).

*In vitro* transcribed tRNAs were used as controls for probe specificity. For the *S. malanga* 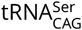 probe, an *in vitro* transcribed version of the target 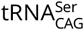 was used as a positive control and an *in vitro* transcribed version of 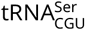 was used as a control for cross-hybridization. For the 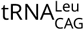 probe, an *in vitro* transcribed version of the target 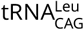 was used as a positive control and an *in vitro* transcribed version of 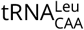 was used as a control for cross-hybridization. Cross-hybridization controls were selected by aligning the reverse complement of the probe sequence using MAFFT v7.429 (***Katoh and Standley, 2013***) with default settings to all tRNA genes in the *S. malanga* genome (found by tRNAscan-SE 2.0), and selecting the non-target tRNA with the highest pairwise alignment score. *In vitro* transcribed tRNAs were produced using the MAXIscript T7 Transcription Kit (Thermo) from a DNA template composed of a T7 promoter (5’ GATCTAATAC-GACTCACTATAGGGAGA 3’) followed by the tRNA sequence. The resulting tRNA transcript has an additional six nucleotides of the promoter included at the 5’ end. CCA-tails were not included in the *in vitro* transcribed tRNA sequences.

Total RNA and *in vitro* transcribed controls for probe specificity were denatured in formamide buffer (Gel Loading Buffer II, Thermo) at 90°C for 5 minutes and electrophoretically separated on a 10% TBE urea gel (Novex). Gels were rinsed in 0.5x TBE and RNA was transferred onto a Hybond N+ membrane (GE Healthcare) in 0.5x TBE by semi-dry transfer (Bio-Rad Transblot) at 3 mA/cm2 for 1 hr. Blots were crosslinked on each side using a Stratalinker UV crosslinker on the “auto-crosslink” setting. Blots were prehybridized in PerfectHyb Plus Hybridization buffer (Sigma) at 64°C for 1 hour prior to incubation with the radiolabelled DNA probe overnight. Blots were washed at 64°C twice in low stringency buffer (0.1% SDS, 2x SSC) for 15 minutes and once in high stringency buffer (0.1% SDS, 0.1x SSC) for 10 minutes, exposed on storage phosphor screens, and scanned using a Typhoon imager.

### Acid urea PAGE Northern blotting for tRNA charging

For the partial deacylation control, total RNA was treated in 100 mM Tris pH 7.0 at 37°C for 30 minutes, quenched with an equal volume of buffer containing 50 mM NaOAc and 100mM NaCl, and precipitated. Electrophoresis on acid urea polyacrylamide gels was performed as described in ***Varshney et al.*** (***1991***). 4 *μ*g of total RNA and partial deacylation control in acid urea sample buffer (0.1 NaOAc pH 4.5, 8M urea, 0.05% bromophenol blue, 0.05% xylene cyanol) were loaded onto a 0.4mm thick 6.5% polyacrylamide gel (SequaGel) containing 8M urea and 100mM NaOAc pH 4.5 and run for 18 hours at 450 V in 4°C with 100mM NaOAc pH 4.5 running buffer. The region between the two dyes corresponds to the tRNA size range, and was cut out and transferred onto a blot for probing following the same steps as above for Northern blotting.

## Acknowledgments

We thank members of the Eddy lab for discussions and for comments on the manuscript, A. Murray and A. Darnell for advice and guidance on Northern blotting experiments, and R. Helmiss for feedback on data presentation. Computations were performed on the Cannon cluster, supported by the Harvard FAS Division of Science’s Research Computing Group. Research reported in this publication was supported by the Howard Hughes Medical Institute and by the National Human Genome Research Institute of the National Institutes of Health under award numbers F31-HG010984 (to YS) and R01-HG009116 (to SRE). The content is solely the responsibility of the authors and does not necessarily represent the official views of the National Institutes of Health.

---

**Figure 2-Figure supplement 1.**
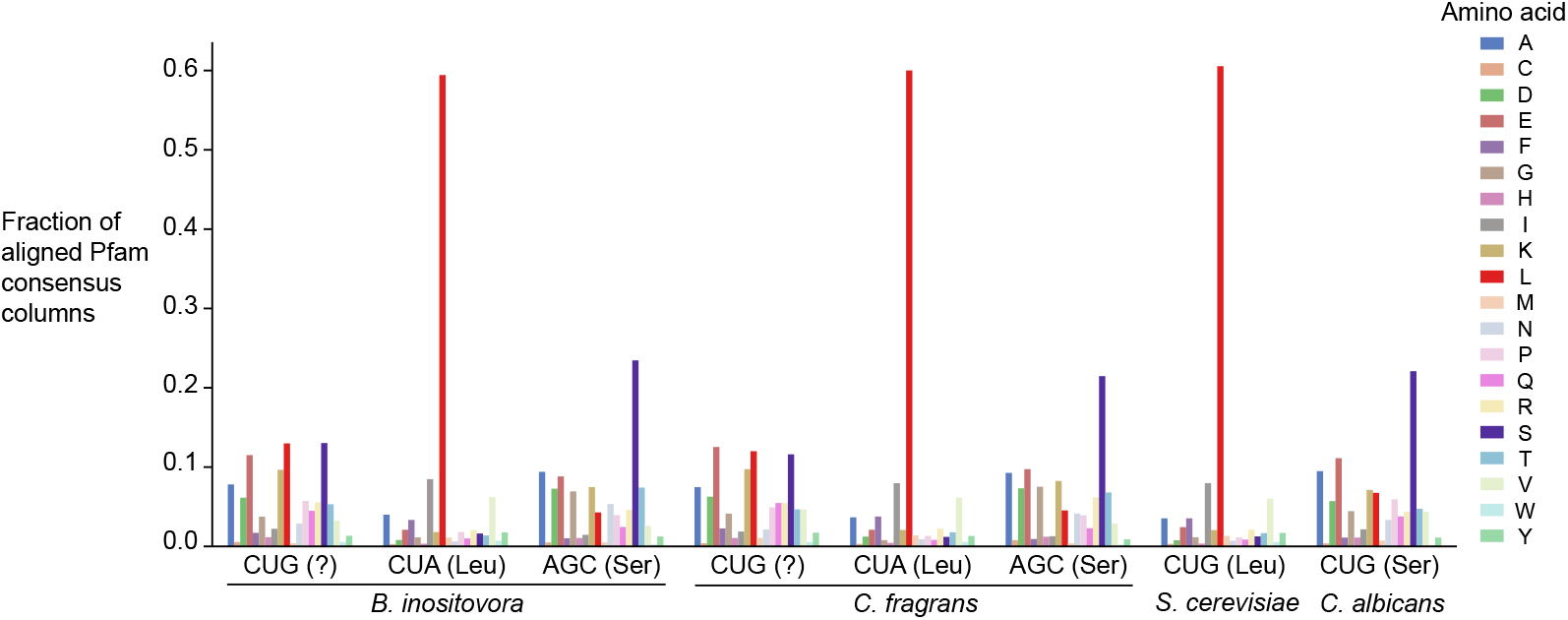
Distribution of the highest probability amino acid for all aligned Pfam consensus columns to CUG, CUA (rare leucine codon), and AGC (rare serine codon) in *B. inositovora* and *C. fragrans* and to CUG in *S. cerevisiae* and *C. albicans*. Codetta codon inference is labelled in parentheses.

**Figure 4-Figure supplement 1.**
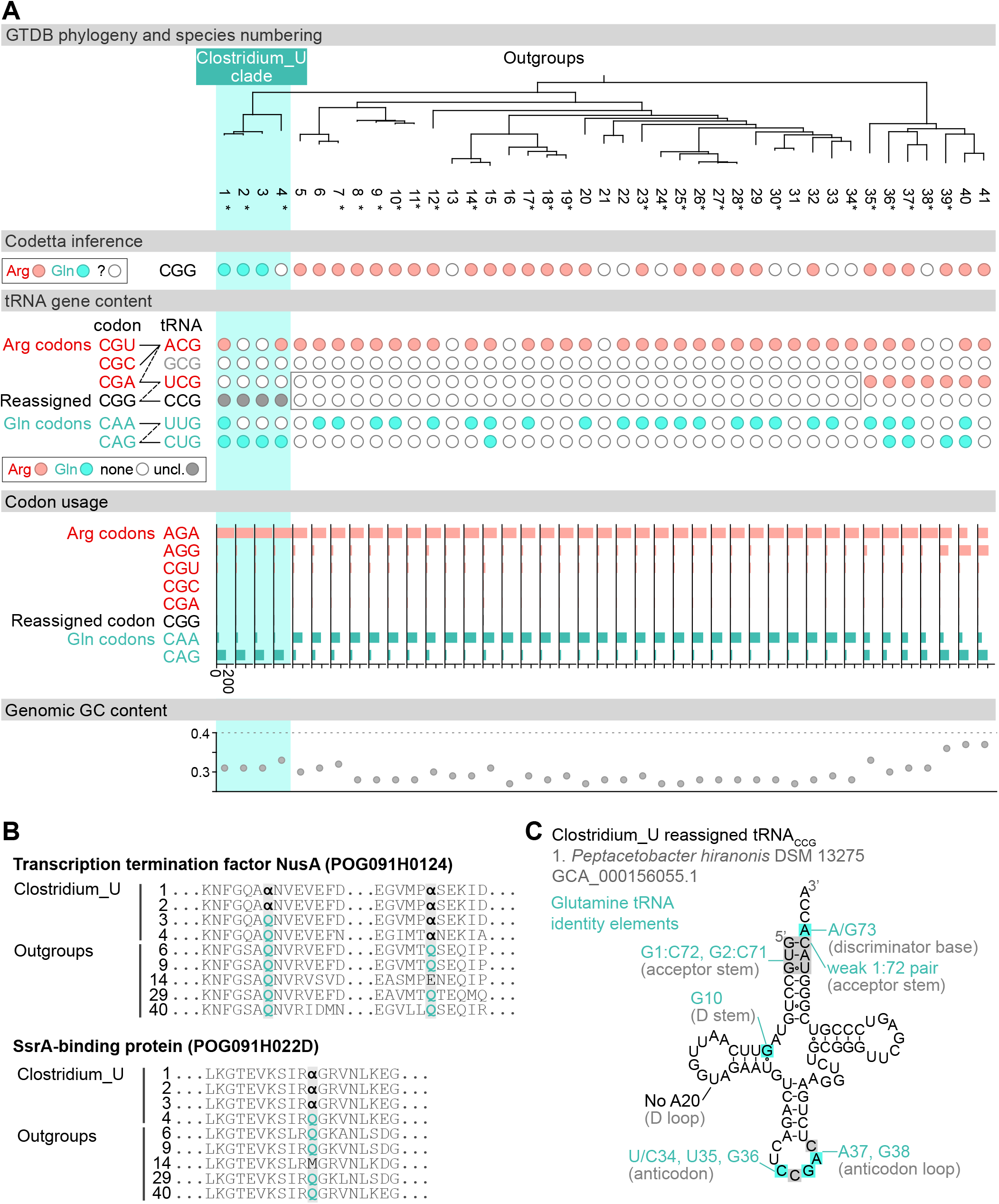
(A) GTDB phylogenetic tree of *Peptacetobacter* (Clostridium_U in GTDB) and closest outgroup genomes. Species numbers can be cross-referenced with ***Figure 4***– ***source data 1***. We consider the entire Clostridium_U clade to have reassigned CGG to glutamine due to the presence of an almost identical CGG-decoding tRNA_CCG_ in all four species. Asterisks indicate genomes with GTDB CheckM estimated genome completeness >99%. For each species, the Codetta CGG inference is indicated by colored circles (red: arginine, light blue: glutamine, white: ‘?’). The presence of tRNA genes that recognize the CAR- and CGN-codons is indicated by filled circles, colored according to the predicted amino acid charging based on based on identity elements for tRNAs (see Methods). A gray box outlines the inability to locate any CGG-decoding tRNAs in the Peptostreptococcaceae (species #5-34). The lines connecting codons and tRNA anticodons represent the likely wobble decoding capabilities, with dashed lines representing weaker interactions. The anticodon ACG is presumed to be modified to ICG, and the U34 of UCG is presumed to be modified in a way that restricts wobble to CGA and CGG, but could potentially recognize CGU and/or CGC depending on the true modification state. Codon usage is the frequency per 10,000 codons aligned to Pfam domains. (B) Multiple sequence alignments of transcription termination factor NusA (BUSCO POG091H0124) and SsrA-binding protein (BUSCO POG091H022D) from the Clostridium_U clade and selected outgroup species. Alignment regions containing nearby CGG (***α***) positions are shown, with columns with CGG in Clostridium_U sequences highlighted. (C) The CGG-decoding tRNA_CCG_ from species #1 (*Peptacetobacter hiranonis* DSM 13275, GCA_000156055.1). tRNA sequence features involved in glutamine identity are highlighted (***Jahn et al., 1991***; ***Hayase et al., 1992***), with nucleotide numbering following the convention of ***Sprinzl et al.*** (***1998***). Nucleotides highlighted in gray do not match the expected identity element.

**Figure 4-Figure supplement 2.**
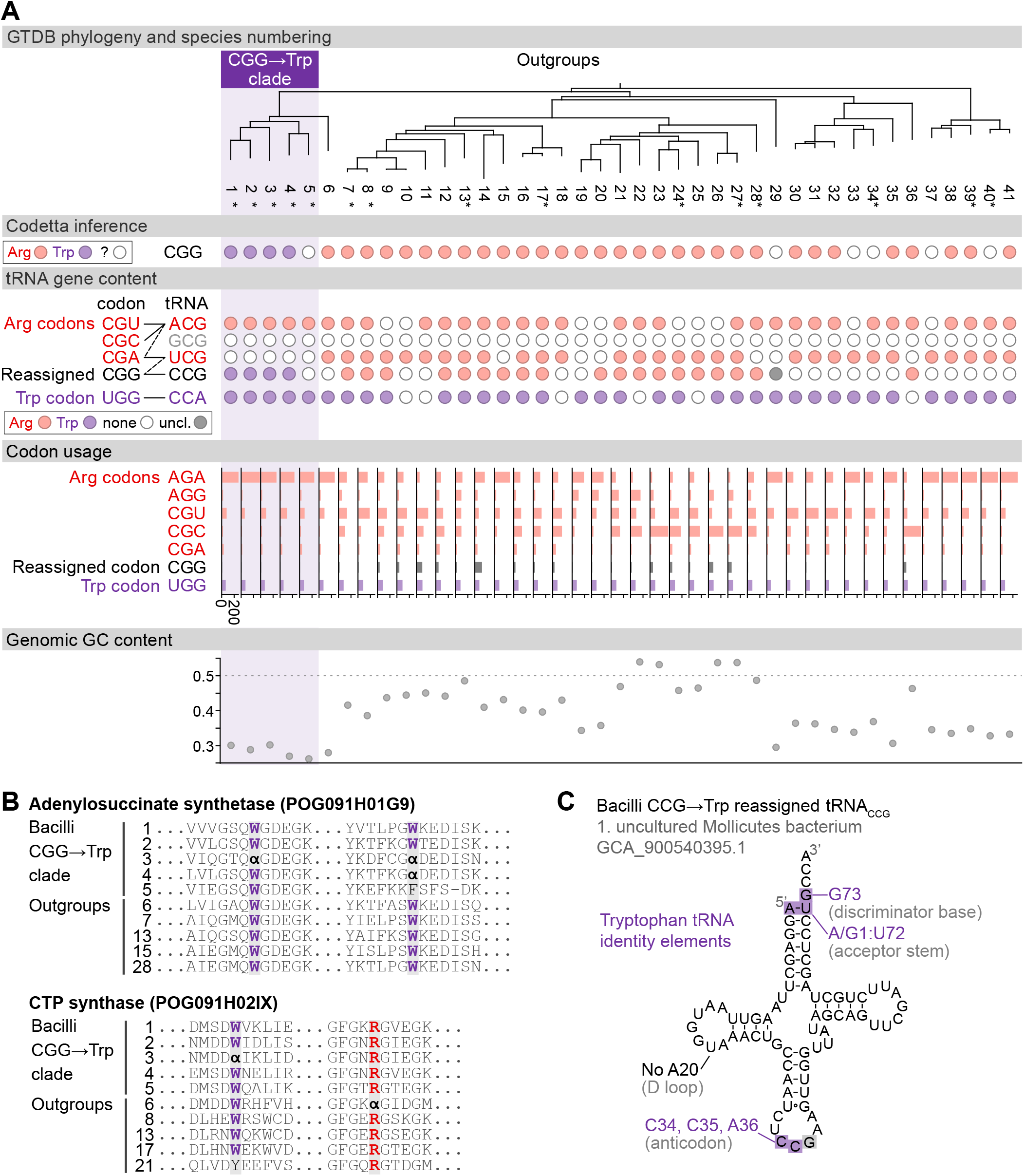
(A) GTDB phylogenetic tree of the Bacilli CGG→Trp clade and closest outgroup genomes. Species numbers can be cross-referenced with ***Figure 4***–***source data 1***. We consider species #5 to be part of the reassigned clade due to the tree topology. Asterisks indicate genomes with GTDB CheckM estimated genome completeness >95%. For each species, the Codetta CGG inference is indicated by colored circles (red: arginine, purple: tryptophan, white: uninferred). The presence of tRNA genes that recognize the UGG and CGN-codons is indicated by filled circles, colored according to the predicted amino acid charging based on identity elements for tRNAs (see Methods). The lines connecting codons and tRNA anticodons represent the likely wobble decoding capabilities, with dashed lines representing weaker interactions. The anticodon ACG is presumed to be modified to ICG. The U34 of anticodon UCG is presumed to be modified in a way that restricts decoding to CGA and CGG, but could potentially recognize CGU and/or CGC depending on the true modification state. Codon usage is the frequency per 10,000 codons aligned to Pfam domains. (B) Multiple sequence alignments of adenylosuccinate synthetase (BUSCO POG091H01G9) and CTP synthase (BUSCO POG091H02IX) from the Bacilli CGG→Trp clade and selected outgroup species. Alignment regions containing CGG (***α***) at conserved positions are shown, with columns with CGG in Bacilli CGG→Trp clade and the closest outgroup (species #6) sequences highlighted. (C) The CGG-decoding tRNA_CCG_ from species #1 (uncultured Mollicutes bacterium, GCA_900540395.1). tRNA sequence features involved in tryptophan identity are highlighted ***(Giegé et al., 1998;Himeno et al., 1991)***, with nucleotide numbering following the convention of ***Sprinzl et al.*** (***1998***). Nucleotides highlighted in gray do not match the expected identity element.

**Figure 4-Figure supplement 3.**
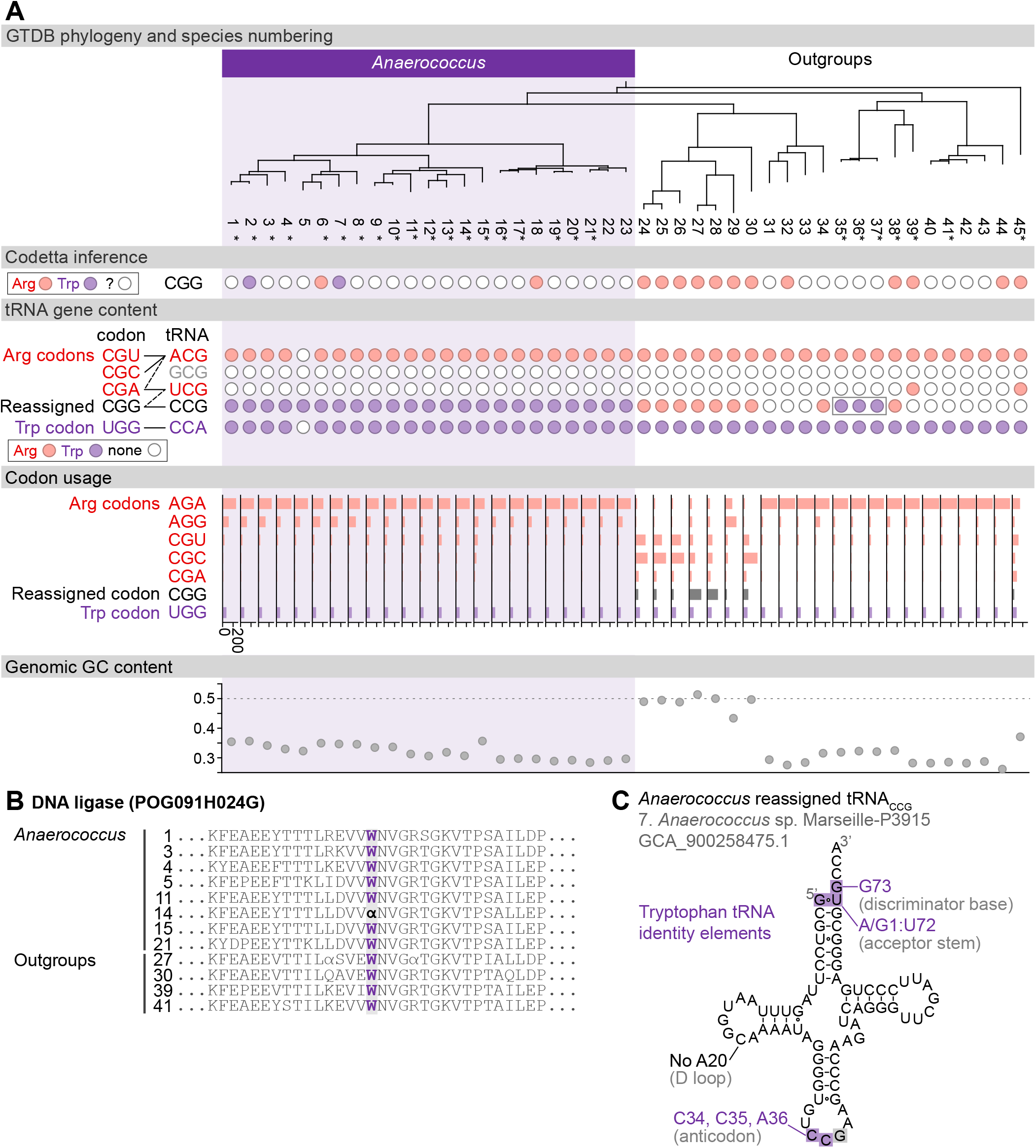
(A) GTDB phylogenetic tree of *Anaerococcus* and closest outgroup genomes. Species numbers can be cross-referenced with ***Figure 4***–***source data 1***. We considered the entire *Anaeroccocus* clade to have reassigned CGG to tryptophan due to the presence of a tryptophan-like tRNA_CCG_ in all *Anaerococcus* species. Asterisks indicate genomes with GTDB CheckM estimated genome completeness >98%. For each species, the translation of the reassigned codon CGG inferred by Codetta is indicated by colored circles (red: arginine, purple: tryptophan, white: ‘?’). The presence of tRNA genes that recognize the UGG and CGN-codons is also indicated by filled circles, colored according to the predicted amino acid charging based on identity elements for tRNAs (see Methods). A gray box outlines the tRNA_CCG_ in *Finegoldia*, which has features of tryptophan identity. The lines connecting codons and tRNA anticodons represent the likely wobble decoding capabilities, with dashed lines representing weaker interactions. The anticodon ACG is presumed to be modified to ICG. The U34 of anticodon UCG is presumed to be modified in a way that restricts decoding to CGA and CGG, but could potentially recognize CGU and/or CGC depending on the true modification state. Codon usage is the frequency per 10,000 codons aligned to Pfam domains. (B) Region of a multiple sequence alignment of DNA ligase (BUSCO POG091H024G) from *Anaerococcus* species and selected outgroup species, containing a CGG (***α***) at a conserved position in a single *Anaerococcus* species. (C) The CGG-decoding tRNA_CCG_ from species #7 (*Anaerococcus* sp. Marseille-P3915, GCA_900258475.1). tRNA sequence features involved in tryptophan identity are highlighted ***(Giegé et al., 1998;Himeno et al., 1991)***, with nucleotide numbering following the convention of ***Sprinzl et al.*** (***1998***). Nucleotides highlighted in gray do not match the expected identity element.

**Figure 4-Figure supplement 4.**
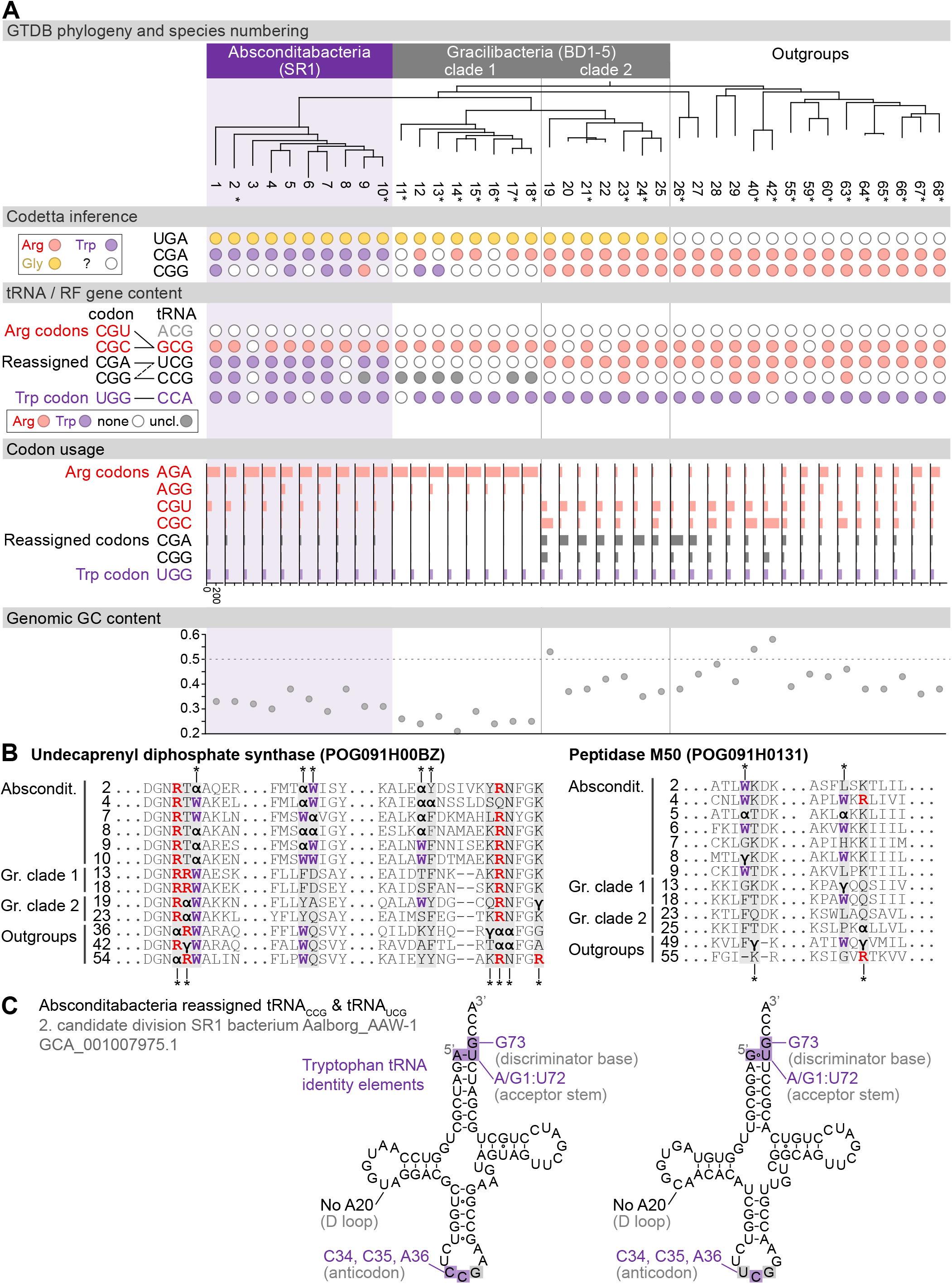
(A) GTDB phylogenetic tree of Absconditabacteria, Gracilibacteria, and closest outgroup genomes. Species numbers can be cross-referenced with ***Figure 4***–***source data 1***. We considered the entire Absconditabacteria clade to have reassigned CGA and CGG to tryptophan due to a combination of Codetta inference, phylogeny, and evidence from tRNA genes and multiple sequence alignments of BUSCO genes. We provisionally split the Gracilibacteria into two clades based on differences in Codetta CGG inference, tRNA gene content, codon usage, and GC content. Gracilibacteria clade 1 may have reassigned CGG to tryptophan, pending additional evidence. Asterisks indicate genomes with GTDB CheckM estimated genome completeness >75%. For each species, the Codetta inference of the three reassigned codons (UGA, CGA, and CGG) is indicated by colored circles (red: arginine, purple: tryptophan, yellow: glycine, white: ‘?’). The presence of tRNA genes that recognize UGG and CGN-codons is indicated by filled circles, colored according to the predicted amino acid charging based on identity elements for tRNAs (see Methods). The lines connecting codons and tRNA anticodons represent the likely wobble decoding capabilities, with dashed lines representing weaker interactions. The U34 of UCG is presumed to be modified in a way that restricts wobble to CGA and CGG, but could potentially recognize CGU and/or CGC depending on the true modification state. Codon usage is the frequency per 10,000 codons aligned to Pfam domains. (B) Multiple sequence alignments of undecaprenyl diphosphate synthase (BUSCO POG091H00BZ) and Peptidase M50 (BUSCO POG091H0131) from Absconditabacteria, Gracilibacteria clades 1 and 2, and selected outgroup species. Alignment regions containing nearby CGA (***α***) or CGG (***γ***) positions are shown, with columns containing CGA or CGG in Absconditabacteria or Gracilibacteria clade 1 sequences highlighted with an asterisk above, and columns containing CGA or CGG in Gracilibacteria clade 2 and outgroup sequences highlighted with an asterisk below. (C) The CGA- and CGG-decoding tRNAs (UCG and CCG anticodons) from species #2 (candidate division SR1 bacterium Aalborg_AAW-1, GCA_001007975.1). tRNA sequence features involved in tryptophan identity are highlighted (***Giegé et al., 1998***; ***Himeno et al., 1991***), with nucleotide numbering following the convention of ***Sprinzl et al.*** (***1998***). Nucleotides highlighted in gray do not match the expected identity element.

